# Lower degree of microsatellite instability in colorectal carcinomas from *MSH6*-associated Lynch syndrome patients

**DOI:** 10.1101/2024.08.12.607570

**Authors:** Noah C. Helderman, Fabian Strobel, Lena Bohaumilitzky, Diantha Terlouw, Anne-Sophie van der Werf – ′t Lam, Tom van Wezel, Hans Morreau, Magnus von Knebel Doeberitz, Maartje Nielsen, Matthias Kloor, Aysel Ahadova

## Abstract

**Background:** Numerous observational and molecular studies focusing on Lynch syndrome (LS) have revealed significant variation in the phenotype and molecular characteristics among carriers of pathogenic variants in mismatch repair genes (*path_MMR*). Recently, we demonstrated that colorectal carcinomas in *path_MSH6* carriers exhibit fewer insertion/deletion mutations compared to CRCs from other MMR groups, raising the question of whether *MSH6*-mutated CRCs might display a lower degree of microsatellite instability (MSI).

**Methods:** Mutations at twenty coding microsatellites (cMS) were analyzed in 39 *MSH6*-, 18 *MLH1*-, 16 *MSH2*- and 22 *PMS2*-mutated CRCs and 35 sporadic MSI CRCs, and mutation frequencies and mutant allele ratios were compared among the different MMR-deficient groups. Considering factors such as *HLA-A*\*02:01 type, *B2M* status, and the anticipated immunogenicity of frameshift peptides derived from cMS mutations, the identified cMS mutation profiles of *MSH6*-mutated CRCs were further investigated to assess their potential impact on immunotherapeutic strategies.

**Results:** *MSH6*-mutated CRCs exhibited lower mutation frequencies and mutant allele ratios across most cMS. The cMS mutations in *MSH6*-mutated CRCs demonstrated inverse correlations with the predicted immunogenicity of the resulting frameshift peptides, which may suggest negative selection of cell clones bearing highly immunogenic frameshift peptides.

**Conclusions:** *MSH6*-mutated CRCs display a lower degree of MSI, which may be connected to lower penetrance and later onset of the disease in *path_MSH6* carriers. Moreover, this lower MSI level may implicate an altered immune response compared to other MSI CRCs, which could have implications for the success of immunotherapy in *MSH6*-mutated CRCs. Future studies should carefully evaluate this possibility. If confirmed, these results would reinforce the notion of classifying LS as distinct syndromes associated with specific MMR genes.

## Introduction

Lynch syndrome (LS) is one of the most common hereditary cancer syndromes, resulting from a germline pathogenic or likely pathogenic variant in one of the mismatch repair genes (*path_MMR*)— *MLH*1, *MSH2*, *MSH6*, or *PMS2*—or, less frequently, from an *EPCAM* deletion located upstream of *MSH2*.^1^

While traditionally considered a single hereditary cancer syndrome, LS has been found to demonstrate significant variability in cumulative colorectal cancer (CRC) risks depending on the affected MMR gene^2–5^, as well as distinct molecular characteristics in the respective CRCs^1, 6–9^, prompting the question of whether LS should be redefined as multiple gene-specific syndromes.^10, 11^ In line with these observations, our recent study demonstrated fewer MMR-deficiency (MMR-d) signature-associated insertion/deletion mutations (INDELs) in CRCs from *path_MSH6* carriers compared to other LS CRCs.^9^ This difference might be attributed to the specific function of MSH6 in DNA MMR, being essential for repairing single nucleotide variants (SNVs) in complex with MSH2 (hMutSα), but being redundant in repairing INDELs, which predominantly rely on the hMutSβ (MSH2-MSH3) complex.^9^

By shifting of the reading frame, INDELs affecting coding microsatellites (cMS) can inactivate tumor suppressor genes and produce frameshift peptides (FSPs).^12^ As these are highly immunogenic to the host’s immune system, microsatellite instable (MSI) CRCs generally trigger robust immune responses and respond well to immune checkpoint blockade therapy.^13–15^ As certain cMS mutations in MSI CRCs are positively selected due to their growth-promoting effect, the resulting FSPs form the basis of a novel vaccine strategy that aims to prevent tumor formation in LS carriers.^16^ Such FSP neoantigen-based vaccines were already proven effective in a LS mouse model^17^ and evoked humoral and cellular immune responses without causing serious adverse events in a phase I/IIa clinical trial (NCT01461148).^18^

Given the potential significance of INDELs/MSI for immunotherapy and preventive vaccine strategies, further investigation into the (immunogenic) cMS mutation pattern of LS CRCs is warranted and may have important implications for *path_MSH6* carriers specifically. Additionally, such research may shed light on the different pathways of colorectal carcinogenesis in *path_MMR* carriers^6, 19^, as well as on the phenotypical differences observed among *path_MMR* carriers, including lower cumulative risks of CRCs for *path_MSH6* (8-17%) and *path_PMS2* (6-11%) carriers compared to *path_MLH1* (27-52%) and *path_MSH2* (23-50%) carriers.^2–4, 20, 21^

Previously, we evaluated and compared immunogenic cMS mutation patterns between *PMS2-* and *MLH1*-/*MSH2*-mutated CRCs, which appeared very similar.^22^ Moreover, recent studies suggest a potential inverse relationship between cMS mutation frequencies and the immunogenicity of cMS mutation-derived FSP epitopes, indicating immunoediting of MSI tumor cell clones.^23, 24^ However, cMS profiles of *MSH6*-mutated CRCs have not been analyzed previously. In the current study, we therefore aimed to define the cMS mutation pattern of *MSH6*-mutated CRCs, compare it with that of other LS CRCs, and analyze the potential consequences of the (HLA-specific) immunogenicity of the cMS mutation-derived FSPs on the cMS mutation patterns in *MSH6*-mutated CRCs.

## Methods

### Ethical statement

This study was approved by the Medical Ethical Committee of Leiden, The Hague, Delft (protocol P17.098). Patient samples were handled according to the medical ethical guidelines described in the Code of Conduct for responsible use of human tissue in the context of health research (Federation of Dutch Medical Scientific Societies). Patients provided informed consent for the use of tissue and data.

### Patients and samples

Coded or anonymized formalin-fixed, paraffin-embedded (FFPE) tumor tissue blocks were obtained from the department of Pathology of the Leiden University Medical Centre (Leiden, The Netherlands) or via the Dutch Pathology Registry (PALGA; reference LZV2022-68), respectively. Collected tumor tissue originated from 39 *MSH6-* CRCs, 18 *MLH1-* CRCs, 16 *MSH2-* CRCs, 22 *PMS2-*mutated CRCs and from 35 sporadic CRCs which were MMR-d due to *MLH1* promotor hypermethylation (*MLH1*-PM). Available clinical and histological data were extracted from pathology reports and/or patient records.

### Tumor workup and DNA isolation

For providers and relevant (user) details of all materials used in this study, see **Supplementary Table 1**. FFPE tissue blocks were cut into 2-4 μm sections and placed on silane-coated glass slides. Sections were deparaffinized and stained with hematoxylin and eosin, as described previously.^25^ Following manual microdissection of normal and tumor tissue separately, DNA was isolated using the Promega ReliaPrep FFPE gDNA Miniprep system (Promega Corporation, Fitchburg, WI, USA) according to the manufacturer’s instructions.

### Immunohistochemical staining for MMR

In cases where MMR status was not reported in available pathology reports and/or patient records, immunohistochemical detection of MMR protein expression was performed following the previously described method and the antibodies listed in **Supplementary Table 1**.^25^

### MSI analysis

In cases, for which MSI status was not reported in available pathology reports and/or patient records, CRCs were characterized for their MSI status by (multiplex) PCR and fragment length analysis of four markers, including BAT25, BAT26, CAT25 and BAT40, as described previously.^26, 27^ CRCs displaying instability in ≥2 of the analyzed markers were classified as MSI.

### cMS analysis

Fragments length analysis using fluorescently labeled primers specific for 20 cMS was performed for all MSI CRCs as previously described.^22, 23^ The selection of cMS was based on a genome-wide search for cMS^28^, and data on likely functional relevance in LS CRCs derived from the publications by Ballhausen et al.^23^, Kloor et al.^18^, and Bajwa-Ten Broeke et al (**Supplementary Table 1**).^22^ PCR fragments were visualized on an ABI3130xl genetic analyzer and generated raw data were analyzed using GeneMarker® Software (v1.5). Peak height profiles were analyzed using ReFrame in RStudio (v.2022.02.01).^23^ cMS with a ≥15% prevalence of non-wild type alleles in a certain sample were classified as mutant. The normal mucosa of nine samples were analyzed as a reference.

Additional cMS data were retrieved from a distinct cohort of 41 *MLH1-*, 21 *MSH2-,* and 12 *PMS2-*mutated CRCs that were previously analyzed and described following a similar protocol.^22^

### *HLA-A*\*02:01 and *B2M* genotyping

*MSH6*-mutated CRCs were genotyped for the *B2M* status following the previously described method and primers listed in **Supplementary Table 1**^23, 29^, and for the *HLA-A*\*02:01 status using a previously described SNP analysis on DNA isolated from FFPE tumor tissue.^23, 24^ This SNP, located in the 5’ region of exon 2 of the *HLA-A* gene, allows highly accurate classification of samples as *HLA-A*\*02:01 positive or negative.^30^ PCR amplification of a region from the intron between exon 1 and 2 to the 5’ region of exon 2 of the *HLA-A* locus was performed using the forward and reverse primers from **Supplementary Table 1**. Subsequently, Sanger sequencing of the reverse (G-R) primer was conducted on an ABI3130xl genetic analyzer^31^ and sequencing data was analyzed using Sequencing Analysis Software. Nucleotide position 78 was used for classification, where a T (homo-/heterozygous) indicated *HLA-A*\*02:01 positivity, while a C (homozygous) at this position indicated *HLA-A*\*02:01 negativity (**Supplementary** Figure 1).

### Immunogenicity of cMS mutation-derived FSPs

To investigate the relationship between cMS mutation frequencies and the immunogenicity of cMS mutation-derived FSP epitopes, we followed previously described methods, considering patients’ *B2M* and *HLA-A*\*02:01 status.^23, 24^

First, the mean mutation frequency and mAR of M1-frameshift mutations was calculated for each cMS. M1-frameshift mutations, denoted as “M1 = minus 1,” involve a reading frame shift of one position to the minus side, including deletions of one nucleotide (L1) or insertions of two nucleotides (R2), which are the most common frameshift mutations in MSI CRCs.

Secondly, we extracted the general epitope likelihood score (GELS) from Ballhausen et al.^23^, which computes the number of predicted epitopes in FSPs resulting from M1-frameshift mutations. These GELS account for MHC ligand prediction and the prevalence of the respective HLA allele in European Caucasians, aligning with the population from which our tumor samples originate, and are based on an HLA-binding probability of *p*_binding_ = 50%. Moreover, we extracted the overall ligand likelihoods (OLLs) for each cMS from Witt et al.^24^, which assess the likelihood that at least one peptide derived from the M1-frameshift mutation is presented by the HLA-A molecule in patients who are either *HLA-A*\*02:01 positive or negative.

Next, we correlated the GELS with the mutation frequencies, initially using an HLA type-independent approach and subsequently accounting for *HLA-A*\*02:01 status. Moreover, we analyzed the absolute differences in the mean mARs of M1-frameshift mutations and the corresponding OLLs between the *HLA-A\**02:01-positive and *HLA-A*\*02:01-negative groups, using the Pearson correlation coefficient.

### Statistical analysis

Statistical analysis was performed in RStudio (version v2022.02.3+492, Team R, Integrated Development for R, MA, USA) and IBM SPSS Statistics for Windows 2017 (version 29.0, International Business Machines Corporation, NY, USA). Continuous outcomes are presented as means with standard deviations (SDs) and/or as medians with interquartile ranges (IQRs), whereas categorical outcomes are presented as proportions. Correlations were tested using Pearson correlation coefficient. cMS mutation frequencies (percentage of tumors harboring cMS mutations) were compared between the MMR groups using Chi-square tests, whereas the mARs of each cMS were compared by one-way ANOVA with Dunnett’s test for post-hoc analysis. Outcomes were generally compared between *MSH6*-mutated CRCs and the other MMR groups and raw *P* values were subsequently adjusted for the number of comparisons and outcomes under investigation using Benjamini & Hochberg correction for multiple testing. *P-*values mentioned in this article are two-tailed and considered statistically significant when *P* < .05.

## Results

### Clinical characteristics

A description of clinical characteristics is presented in **Table 1**. Of note, the age at CRC diagnosis of *path_MSH6* carriers was higher as compared to the age at CRC diagnosis of *path_MLH1* (*P*=.046), *path_MSH2* (*P*=.013) and *path_PMS2* (*P*=.005) carriers (**Supplementary** Figure 2). Moreover, *MSH6*-mutated CRCs were generally located more distally (i.e. descending colon or rectosigmoid) compared to CRCs from other groups, with a significant difference between *MSH6*-mutated CRCs, and sporadic CRCs which were MMR-d due to *MLH1* promotor hypermethylation (*MLH1*-PM) (*P*<.001).

**Table 1.** Cohort description.

| Descriptive | *MSH6*<br>(n=39) | *MLH1*<br>(n=18) | *MLH1-PM*<br>(n=35) | *MSH2*<br>(n=16) | *PMS2*<br>(n=22) |
| --- | --- | --- | --- | --- | --- |
| **Sex (male)** | 24 (62%) | 9 (50%) | 6 (17%) | 11 (69%) | 10 (45%) |
| **Age at CRC diagnosis** |  |  |  |  |  |
| Median (IQR) | 65.0 (46.8-71.0) | 48.0 (41.0-60.3) | 66.0 (59.0-71.0) | 51.0 (38.5-58.8) | 46.0 (41.0-57.8) |
| Mean (SD) | 60.5 (13.7) | 50.1 (12.4) | 65.9 (9.4) | 48.3 (13.0) | 48.7 (11.2) |
| NA | 1 (3%) | 0 | 0 | 0 | 0 |
| **Side** |  |  |  |  |  |
| Right<sup>a</sup> | 19 (49%) | 13 (72%) | 32 (91%) | 11 (69%) | 14 (64%) |
| Left<sup>b</sup> | 18 (46%) | 2 (6%) | 2 (6%) | 5 (31%) | 8 (36%) |
| NA | 2 (5%) | 3 (17%) | 1 (3%) | 0 | 0 |
| **Location** |  |  |  |  |  |
| Cecum/ascending colon | 14 (36%) | 11 (61%) | 19 (54%) | 8 (50%) | 10 (45%) |
| Transverse colon | 4 (10%) | 2 (11%) | 12 (34%) | 3 (19%) | 1 (5%) |
| Descending colon | 3 (8%) | 1 (6%) | 1 (3%) | 1 (6%) | 1 (5%) |
| Rectosigmoid | 15 (38%) | 1 (6%) | 1 (3%) | 4 (25%) | 7 (32%) |
| NA | 3 (8%) | 3 (17%) | 2 (6%) | 0 | 3 (14%) |
| **Differentiation** |  |  |  |  |  |
| Well | 3 (8%) | 0 | 0 | 1 (6%) | 0 |
| Well/moderate | 4 (10%) | 4 (22%) | 6 (17%) | 4 (25%) | 6 (27%) |
| Moderate | 19 (48%) | 7 (39%) | 7 (20%) | 7 (44%) | 9 (41%) |
| Moderate/poor | 5 (13%) | 1 (6%) | 6 (17%) | 2 (13%) | 2 (9%) |
| Poor | 4 (10%) | 1 (6%) | 7 (20%) | 0 | 2 (9%) |
| NA | 4 (10%) | 5 (28%) | 9 (26%) | 2 (13%) | 3 (14%) |
| **Diameter [cm]** |  |  |  |  |  |
| Median (IQR) | 5.5 (3.3-7.0) | 3.8 (2.0-5.0) | 4.9 (2.6-7.0) | 2.5 (2.0-6.0) | 5.0 (3.5-6.5) |
| Mean (SD) | 6.5 (7.4) | 5.2 (5.8) | 5.0 (2.4) | 5.5 (7.0) | 6.7 (7.6) |
| NA | 7 (18%) | 3 (17%) | 3 (9%) | 5 (31%) | 5 (23%) |
| **MSI status** |  |  |  |  |  |
| MSI | 38 (97%) | 18 (100%) | 35 (100%) | 15 (94%) | 20 (91%) |
| MSS | 1 (3%) | 0 | 0 | 1 (6%) | 2 (9%) |

<sup>a</sup>Right-sided locations include the cecum/ascending colon and transverse colon.
<sup>b</sup>Left-sided locations include the descending colon, sigmoid colon and rectum
CRC, colorectal cancer; IQR, interquartile range; *MLH1*-PM = *MLH1* promotor hypermethylation; MSI, microsatellite instable; MSS, microsatellite stable; NA, not available/applicable; SD, standard deviation

Comparison of cMS mutation patterns of *MSH6*-mutated CRCs and other MMR mutation groups Except for four MSS CRCs, all tumors from every MMR group were MSI and the majority of tumors lacked expression of the corresponding MMR protein(s) in line with the affected MMR gene (**Supplementary Table 2**). The four MSS CRCs were excluded from further analysis.

To study the cMS mutation patterns of the MSI CRCs from every MMR group, we performed fragment length analysis using fluorescently labeled primers specific for 20 cMS as previously described.^22–24^ The selection of these 20 cMS was guided by a genome-wide search for cMS^28^ and supported by data on potential functional relevance as drivers in LS CRCs and their immunological significance, as reported in studies by Ballhausen et al.^23^, Kloor et al.^18^, and Bajwa-Ten Broeke et al.^22^ For every cMS, we determined the mutation status and calculated the frequency of mutant alleles (mutant allele ratio (mAR)) per tumor, with the latter measure being based on all frameshift peptide reading frames combined. Consistent with existing literature^23, 24^, the M1-frameshift peptide reading frame, which includes deletions of one nucleotide (L1) or insertions of two nucleotides (R2), was the most prevalent frameshift peptide reading frame across all MMR groups (**Figure 1A**).

**Figure 1.**
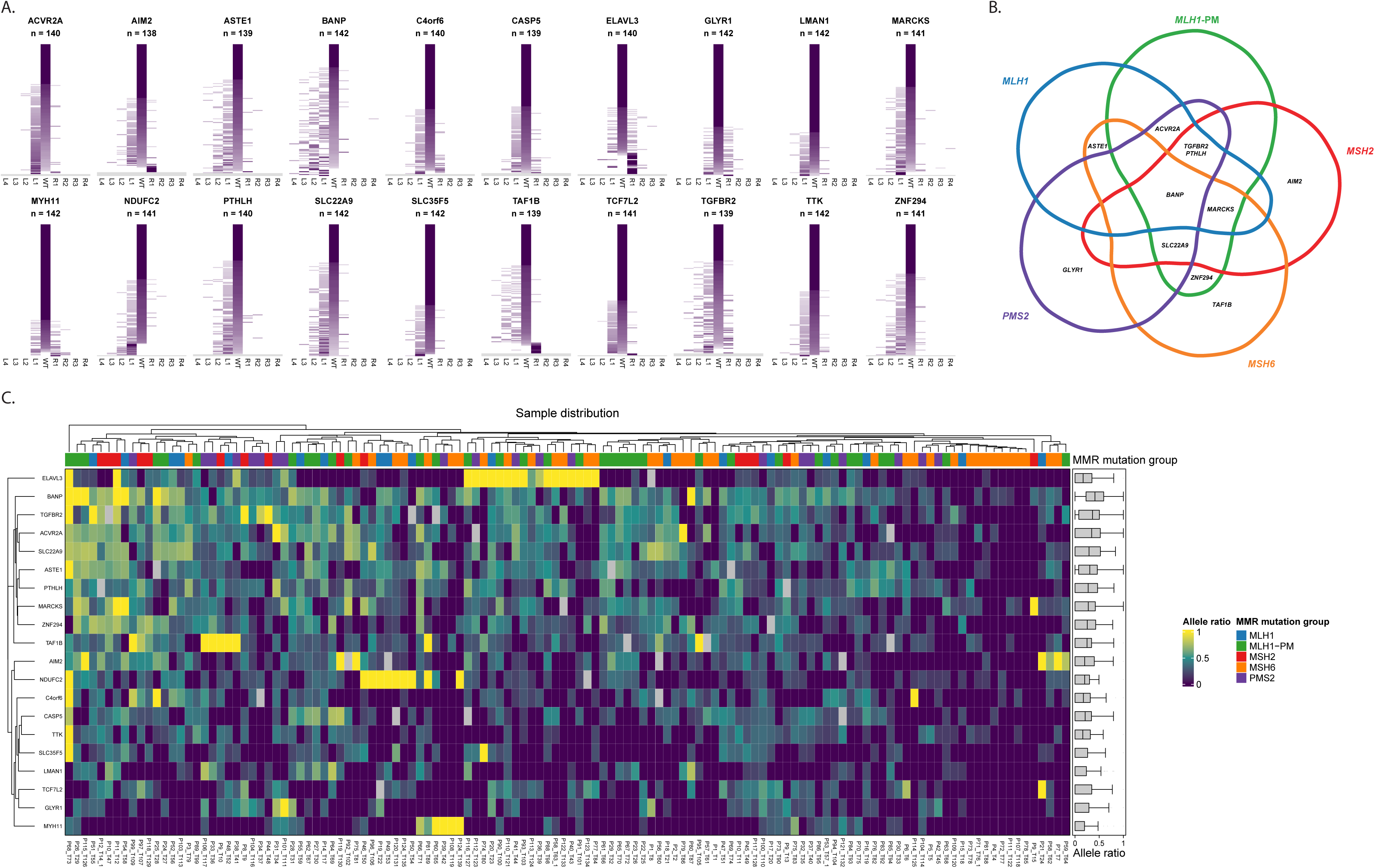
cMS mutation patterns. **A.** Detailed mutational patterns of the 20 analyzed cMS, including mARs for all possible resulting frameshift mutations are depicted. Frameshifts indicated by L represent deletions of 1-4 nucleotides (L1-L4), whereas frameshifts indicated by R represent insertions of 1-4 nucleotides. The intensity of the purple color corresponds with the frequency of the mutant alleles, with lower intensities indicating lower mARs. All MSI CRCs from the current cohort were considered, with each row representing one analyzed CRC. Grey rows represent missing samples. **B.** Venn diagram depicting overlap between the six/seven most frequently mutated cMS per MMR group. **C.** Heatmap depicting unsupervised hierarchic clustering of mARs of the 20 analyzed cMS. All MSI CRCs from the current cohort were considered. *cMS, coding microsatellite; CRC, colorectal cancer; mAR, mutant allele ratio; MLH1-PM, MLH1 promotor hypermethylation; MMR, mismatch repair; MSI, microsatellite instability*.

In *MSH6*-mutated CRCs, *TAF1B* (70%, 26/37), *ASTE1* (68%, 26/38) and *BANP* (68%, 26/38) were most frequently mutated cMS (**Table 2**), with mean mARs of 0.248, 0.239 and 0.264, respectively (**Table 3**). Notably, *BANP* was also among the six most mutated cMS in every other MMR group (**Figure 1B**), exhibiting the highest mean mARs among all cMS in *MLH1-*mutated (mAR 0.504) and *MLH1-*PM (mAR 0.545) CRCs.

**Table 2.**
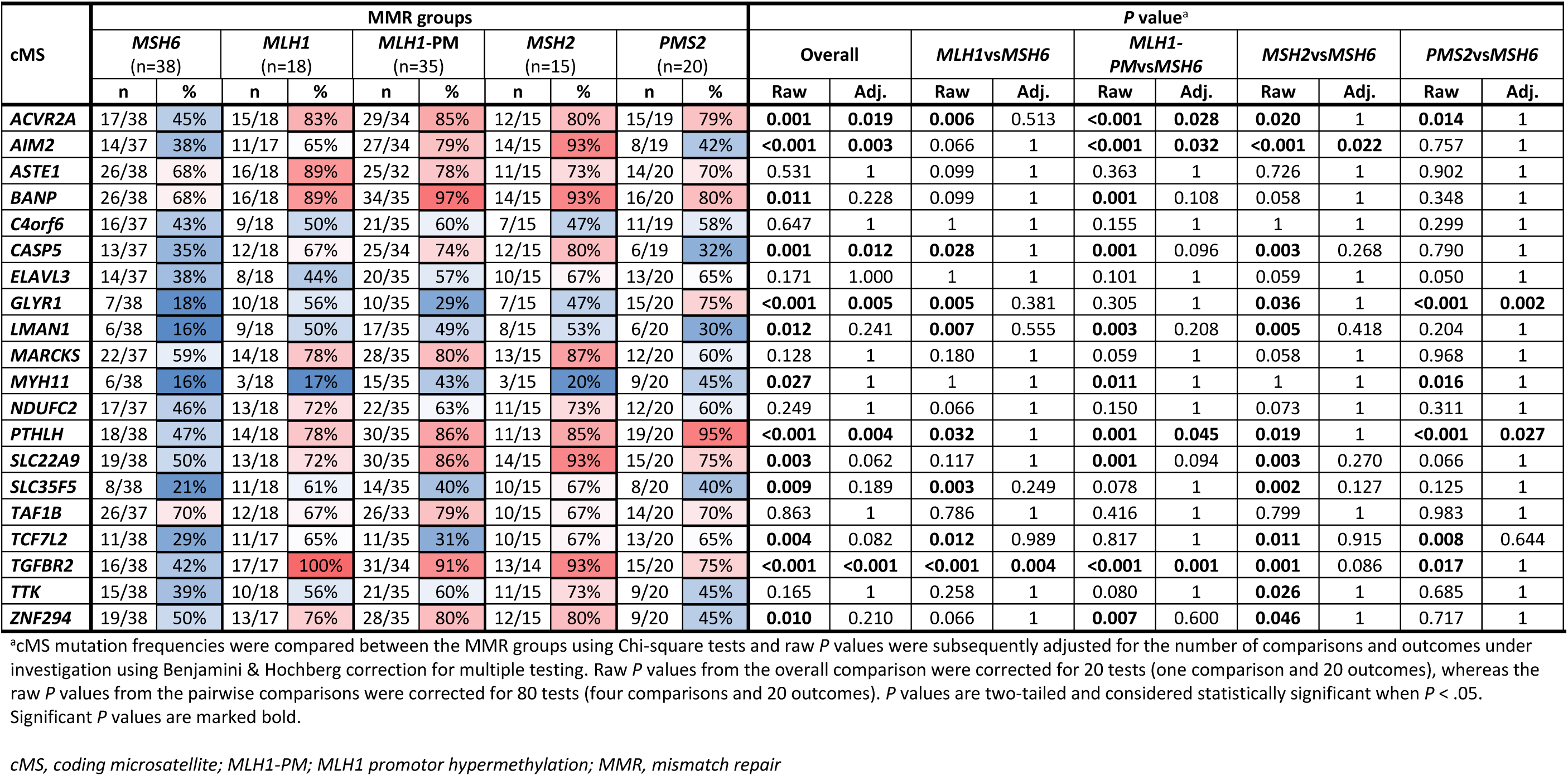
cMS mutation frequencies per MMR group.

**Table 3.** Frequency of mutant cMS alleles per MMR group.

| cMS | mAR per MMR group<sup>a</sup> |  |  |  |  | P value<sup>b</sup> |  |  |  |  |  |  |  |  |  |
| --- | --- | --- | --- | --- | --- | --- | --- | --- | --- | --- | --- | --- | --- | --- | --- |
|  | MSH6<br>(n=38) | MLH1<br>(n=18) | MLH1<br>-PM<br>(n=35) | MSH2<br>(n=15) | PMS2<br>(n=20) | Overall |  | MLH1vsMSH6 |  | MLH1-PMvsMSH6 |  | MSH2vsMSH6 |  | PMS2vsMSH6 |  |
|  |  |  |  |  |  | Raw | Adj. | Raw | Adj. | Raw | Adj. | Raw | Adj. | Raw | Adj. |
| ACVR2A | 0.212 | 0.363 | 0.419 | 0.343 | 0.333 | **0.014** | 0.286 | 0.131 | 1 | **0.003** | 0.051 | 0.287 | 1 | 0.282 | 1 |
| AIM2 | 0.154 | 0.242 | 0.337 | 0.405 | 0.124 | **0.001** | **0.015** | 0.597 | 1 | **0.009** | 0.181 | **0.005** | 0.096 | 0.982 | 1 |
| ASTE1 | 0.239 | 0.361 | 0.39 | 0.273 | 0.239 | **0.028** | 0.563 | 0.192 | 1 | **0.020** | 0.394 | 0.970 | 1 | 1 | 1 |
| BANP | 0.264 | 0.504 | 0.545 | 0.507 | 0.331 | **<0.001** | **<0.001** | **0.003** | 0.054 | **<0.001** | **<0.001** | **0.005** | 0.093 | 0.750 | 1 |
| C4orf6 | 0.187 | 0.207 | 0.225 | 0.165 | 0.214 | 0.526 | 1 | 0.706 | 1 | 0.336 | 1 | 0.988 | 1 | 0.623 | 1 |
| CASP5 | 0.116 | 0.253 | 0.283 | 0.269 | 0.117 | **0.002** | **0.038** | 0.072 | 1 | **0.003** | 0.058 | 0.056 | 1 | 1 | 1 |
| ELAVL3 | 0.261 | 0.183 | 0.298 | 0.221 | 0.369 | 0.413 | 1 | 0.989 | 1 | 0.780 | 1 | 1 | 1 | 0.378 | 1 |
| GLYR1 | 0.058 | 0.187 | 0.104 | 0.175 | 0.342 | **<0.001** | **<0.001** | 0.082 | 1 | 0.749 | 1 | 0.178 | 1 | **<0.001** | **<0.001** |
| LMAN1 | 0.057 | 0.161 | 0.192 | 0.18 | 0.117 | 0.058 | 1 | 0.254 | 1 | **0.021** | 0.428 | 0.173 | 1 | 0.704 | 1 |
| MARCKS | 0.176 | 0.312 | 0.353 | 0.463 | 0.22 | **<0.001** | **0.009** | 0.146 | 1 | **0.006** | 0.116 | **<0.001** | **0.007** | 0.913 | 1 |
| MYH11 | 0.116 | 0.053 | 0.162 | 0.051 | 0.178 | 0.214 | 1 | 0.926 | 1 | 0.572 | 1 | 0.933 | 1 | 0.522 | 1 |
| NDUFC2 | 0.206 | 0.336 | 0.227 | 0.249 | 0.149 | 0.308 | 1 | 0.307 | 1 | 0.994 | 1 | 0.966 | 1 | 0.888 | 1 |
| PTHLH | 0.171 | 0.323 | 0.35 | 0.323 | 0.326 | **0.004** | 0.078 | **0.047** | 0.949 | **0.002** | **0.032** | 0.092 | 1 | **0.032** | 0.648 |
| SLC22A9 | 0.226 | 0.367 | 0.373 | 0.403 | 0.299 | 0.067 | 1 | 0.183 | 1 | 0.056 | 1 | 0.087 | 1 | 0.726 | 1 |
| SLC35F5 | 0.084 | 0.187 | 0.14 | 0.235 | 0.106 | 0.086 | 1 | 0.217 | 1 | 0.594 | 1 | **0.044** | 0.877 | 0.987 | 1 |
| TAF1B | 0.248 | 0.223 | 0.267 | 0.313 | 0.335 | 0.653 | 1 | 0.994 | 1 | 0.996 | 1 | 0.864 | 1 | 0.614 | 1 |
| TCF7L2 | 0.126 | 0.286 | 0.092 | 0.288 | 0.223 | **0.002** | **0.046** | **0.038** | 0.756 | 0.911 | 1 | **0.048** | 0.954 | 0.314 | 1 |
| TGFBR2 | 0.177 | 0.438 | 0.376 | 0.54 | 0.369 | **<0.001** | **<0.001** | **0.001** | **0.018** | **0.002** | **0.040** | **<0.001** | **<0.001** | **0.015** | 0.297 |
| TTK | 0.094 | 0.189 | 0.189 | 0.255 | 0.158 | **0.034** | 0.687 | 0.220 | 1 | 0.089 | 1.000 | **0.014** | 0.289 | 0.545 | 1 |
| ZNF294 | 0.174 | 0.307 | 0.328 | 0.425 | 0.166 | **0.001** | **0.014** | 0.159 | 1 | **0.017** | 0.336 | **0.002** | **0.033** | 1 | 1 |

<sup>a</sup>Tumors that were wildtype and tumors that were mutant for the cMS were both included.
<sup>b</sup>cMS mARs were compared by one-way ANOVA with Dunnett's test for post-hoc analysis. For the pairwise comparisons, raw *P* values were additionally corrected for the number of comparisons (*n*=4) using Benjamini & Hochberg correction for multiple testing. *P* values are two-tailed and considered statistically significant when *P* < .05. Significant *P* values are marked bold.
cMS, coding microsatellite; mAR, mutant allele ratio; MLH1-PM; MLH1 promotor hypermethylation; MMR, mismatch repair

Across most cMS, *MSH6*-mutated CRCs displayed the lowest mutation frequencies and mARs of all MMR groups, also after correction for multiple testing. This distinctive cMS mutation pattern was further reflected by unsupervised hierarchic clustering of the cMS mARs of all tumors, which revealed that a substantial proportion of *MSH6*-mutated CRCs clustered on the side of the heatmap with the lowest cMS mARs (**Figure 1C**).

To further expand the analysis of the comparison of the *MSH6*-mutated CRCs against the other MMR-d CRCs, we performed a combined analysis of the current cohort with a separate cohort previously analyzed and described following a similar protocol.^22^ This combined analysis encompassed an additional 41 *MLH1-*, 21 *MSH2-,* and 12 *PMS2*-mutated CRCs and yielded similar observations (**Supplementary Table 3-4**, **Supplementary** Figure 3). Neither the number of mutated cMS nor the mean cMS mAR per sample correlated with age at CRC diagnosis or with tumor diameter (**Supplementary** Figure 4).

### cMS mutation patterns in different tumor regions and precancerous lesions

To address tumor heterogeneity, the cMS mutations patterns from multiple tumor regions (defined based on histomorphology) were separately analyzed for four CRCs, from which sufficient tumor material from different tumor regions was available (**Figure 2A**, **Supplementary Table 5**). Whilst there was considerable overlap in the cMS mutation patterns of the different tumor regions, they were not identical. In addition to modest differences in the allele ratios of some cMS mutations, a subset of cMS mutations was exclusively present/absent in only one of the tumor regions, irrespective of the underlying MMR gene defect.

**Figure 2.**
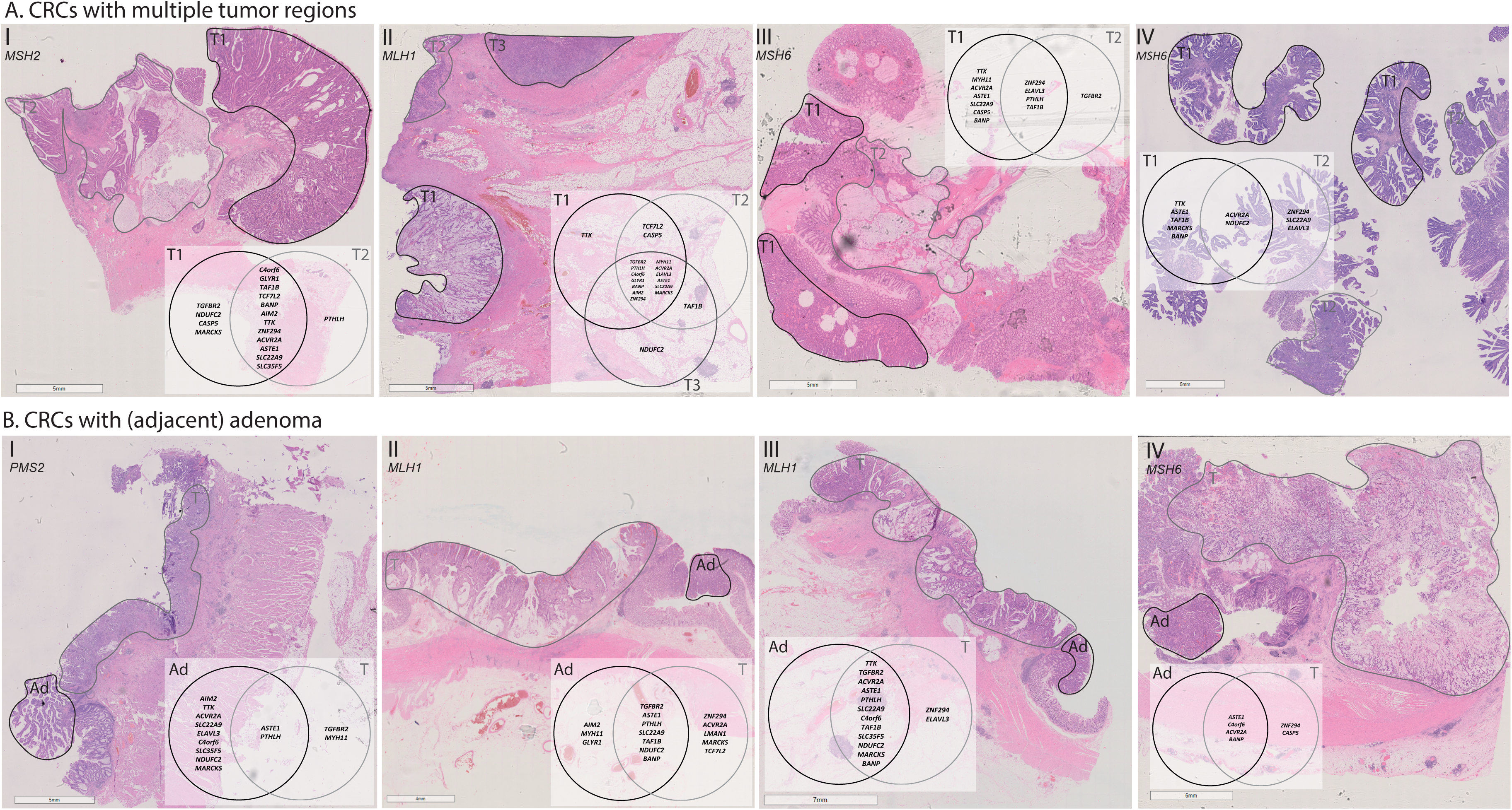
CRCs with multiple tumor regions or (adjacent) adenomas. **A.** The different tumor regions of four CRCs are shown on the respective H&E stains. The Venn diagrams represent the genes containing cMS mutations per tumor region. **B.** Per sample, the adenoma and CRC regions are shown on the respective H&E stains. The Venn diagrams represent the genes containing cMS mutations per tumor/adenoma. IDs correspond with **Supplementary Table 5**. *cMS, coding microsatellite, CRC, colorectal cancer; H&E, hematoxylin and eosin*.

A similar phenomenon was observed when comparing the cMS mutation patterns between four CRCs and adjacent adenomatous tissue, with a subset of cMS mutations being exclusively found in either the CRC or the respective adenomatous tissue (**Figure 2B**, **Supplementary Table 5**). *ZNF294* was mutated in three of the four CRCs but not in the adjacent adenomatous tissues.

### Immunogenicity of cMS mutation-derived FSPs

Recent studies analyzing cMS patterns in MSI cancers have identified a potential inverse correlation between cMS mutation frequencies and the immunogenicity of cMS mutation-derived FSP epitopes.^23, 24^ This suggests possible immunoediting during MSI cancer evolution, where highly immunogenic cMS mutations undergo counterselection in emerging cancer cell clones. To investigate the potential relationship between immunogenicity and mutation frequency specifically in the context of *MSH6* defects, we examined the predicted immunogenicity of cMS mutation-derived FSP epitopes in *MSH6*-mutated CRCs, following previously described methods.^23, 24^

First, we genotyped the *B2M* gene and the *HLA-A*\*02:01 allele, the most common *HLA-A* allele in the Caucasian population, in the *MSH6*-mutated CRCs, as summarized in **Figure 3A**. Next, we correlated the number of predicted epitopes in (M1) cMS mutation-induced FSPs (using the general epitope likelihood score (GELS) adapted from Ballhausen et al.^23^ (**Supplementary Table 8**)) with the frequency of the respective cMS mutations in *MSH6*-mutated CRCs, initially using an HLA type-independent approach. We observed a significant inverse correlation between GELS and mutation frequency when considering very-low or low-affinity HLA-binding, indicating that a high GELS was related to lower mutation frequency (**Figure 3B**; **Supplementary** Figure 5). This implies that there is counterselection of emerging tumor cell clones with highly immunogenic FSPs, suggesting that frameshift peptide/cMS mutation patterns in *MSH6*-mutated CRCs may be influenced by immunoediting. Consistent with prior observations^23^, this significant inverse correlation was only detected among *B2M*-wildtype tumors. In *B2M*-mutant tumors, where immune selection based on HLA class I antigen presentation does not apply, only a trend was observed, possibly reflecting immune surveillance effects prior to the B2M mutation.^23^

**Figure 3.**
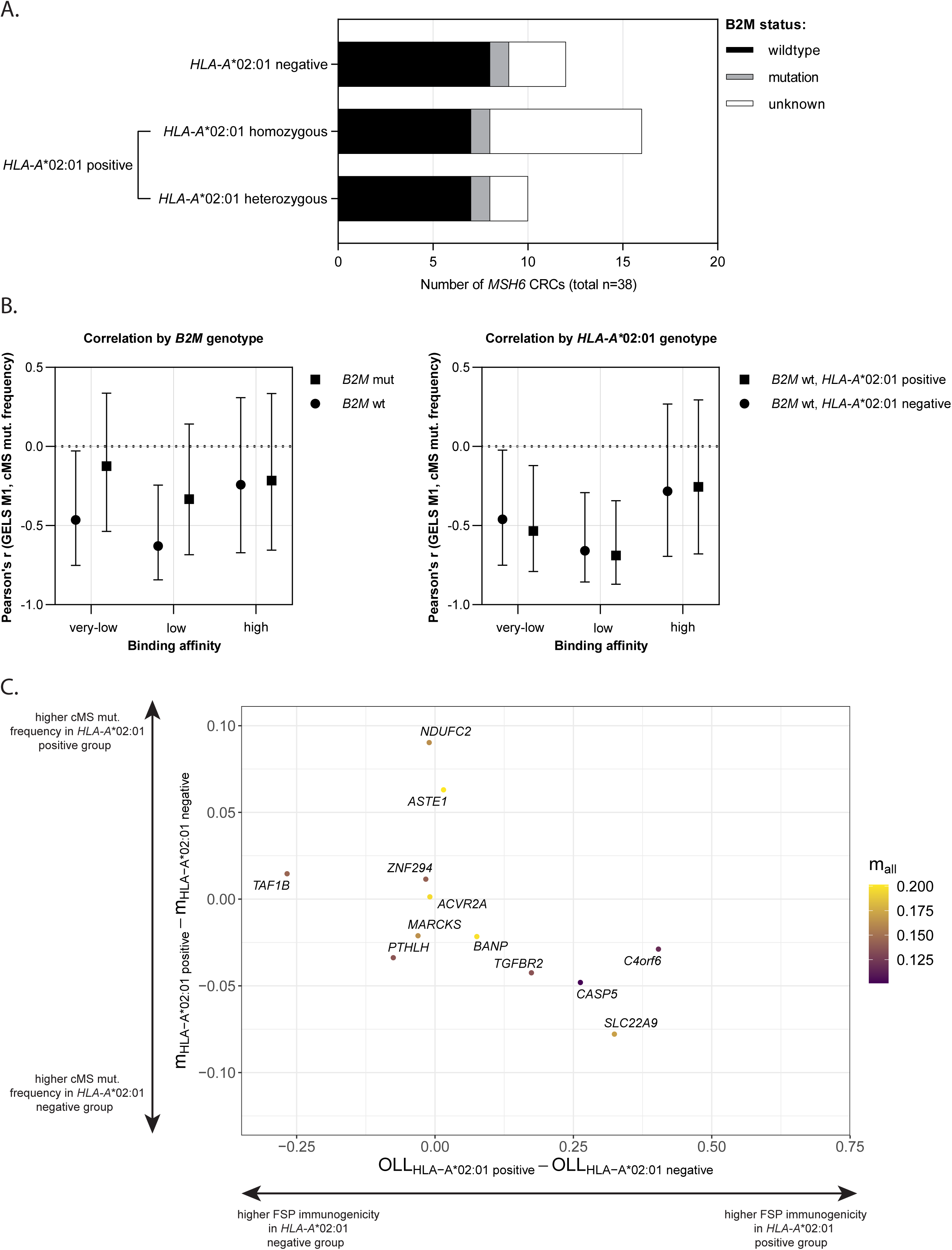
*HLA-A* status of *MSH6*-mutated CRCs and association between cMS mutation frequencies and predicted immunogenicity of cMS mutation-derived FSPs. **A.** The *HLA-A*\*02:01 type and *B2M* status are depicted for all *MSH6*-mutated CRCs (n=38). Samples homo- and heterozygous for the *HLA-A*02:01* allele were both considered to be *HLA-A*\*02:01 positive. **B.** Correlation between the number of predicted epitopes in M1 cMS mutation-derived FSPs (GELS) and the frequency of the respective cMS mutations, categorized by *B2M* mutation status and *HLA-A*\*02:01 status. The y-axis displays Pearson’s r (center points indicating Pearson’s r, with the whiskers representing the 95% confidende intervals), while the x-axis shows the different tumor groups. The GELS, adapted from Ballhausen et al.^23^, are based on MHC ligand prediction and the prevalence of the respective HLA allele in European Caucasians, with a conservative estimate of predicted HLA-binding probability of *p*_binding_ = 50%. **C.** Correlation between the absolute difference of the estimated immunogenicity of cMS mutation-derived FSPs (*x*-axis, OLL_HLA-A*02:01_-OLL_HLA-A*02:01_) and the absolute difference of the underlying M1-frameshift mARs (*y*-axis, m_HLA-A*02:01_-m_HLA-A*02:01_) between the *HLA-A*\*02:01 positive and *HLA-A*\*02:01 negative groups. Only M1-frameshift mutations with a mean mARs > 10% were considered. The intensity of the data points (m_all_) corresponds with the mean M1-frameshift mARs. All *MSH6*-mutated CRCs were included. *cMS, coding microsatellite; CRC, colorectal cancer; FSP, frameshift peptide; GELS, general epitope likelihood score; mAR, mutant allele ratio; OLL, overall ligand likelihood*.

To specifically address the HLA-type dependence of immunoediting, we correlated the GELS and cMS mutation frequencies in the *B2M*-wildtype *HLA-A*\*02:01-positive and *HLA-A*\*02:01-negative groups. Unlike prior observations^23^, we found no HLA class-I restricted effect, as significant negative correlations were observed in both *HLA-A*\*02:01-positive and *HLA-A*\*02:01-negative groups (**Figure 3B**; **Supplementary** Figure 5). However, when correlating the M1-frameshift mARs with the likelihood that at least one peptide derived from the M1-frameshift mutation is presented by the HLA-A molecule in patients who are either *HLA-A*\*02:01-positive or *HLA-A*\*02:01-negative (using the overall ligand likelihoods (OLLs) from Witt et al.^24^ (**Supplementary Table 8**)), we noted an inverse correlation of borderline significance (*P* = .072; Pearson’s r −0.536, 95% CI −0.849 to 0.055), which may support the concept of HLA class I-related immunoediting (**Supplementary** Figure 3C).

## Discussion

An increasing number of epidemiological and molecular studies on LS have highlighted considerable variety in the clinical phenotype and molecular characteristics of *path_MMR* carriers. These findings have raised the question whether LS should still be considered a single hereditary cancer syndrome or multiple syndromes defined by different MMR groups.^10, 11^ We recently showed that CRCs from *path_MSH6* carriers contain fewer MMR-d signature-associated INDELs than CRCs from other MMR groups.^9^ This finding suggests fundamental differences in the pathogenesis of *MSH6*-mutated CRCs and may further indicate the need for subgroup-specific guidelines for screening and treatment.

In the current study, we ought to validate our previous finding by studying the (immunogenic) cMS mutation patterns of LS CRCs. Our new findings, showing that *MSH6*-mutated CRCs generally contain fewer cMS mutations than CRCs from other MMR groups, are in line with our previous observations and could in theory have important molecular and clinical implications.

First, our findings support for the hypothesis that *MSH6*-mutated CRCs generally display a relatively lower degree of MSI. This is likely due to the functional redundancy of MSH6 in INDEL repair (**Figure 4A**), as discussed earlier, which implies that SNVs rather than INDELs drive the development of *MSH6*-mutated CRCs (**Figure 4B**). The reduced accumulation of INDELs/cMS mutations following MSH6 deficiency may, in theory, be associated with a slower tumor progression, a hypothesis which could explain the higher age at CRC diagnosis of *path_MSH6* carriers as observed in our cohort as well as in previous studies.^9, 32–34^ This hypothesis might be controversial, as the reduced accumulation of cMS would also result in lower expression of neoantigens, possibly reducing the immune system’s pressure on the tumor. Given the observed fact that cancer onset occurs at an older age in *path_MSH6* carriers, it likely suggests that the less severe impact of *MSH6* variants – due to the redundant function of the MSH6-MSH2 and MSH3-MSH2 heterodimers in INDEL repair – is more significant for these patients than the decreased immune pressure on MMR-d crypts, precursor lesions, and tumors.

**Figure 4.**
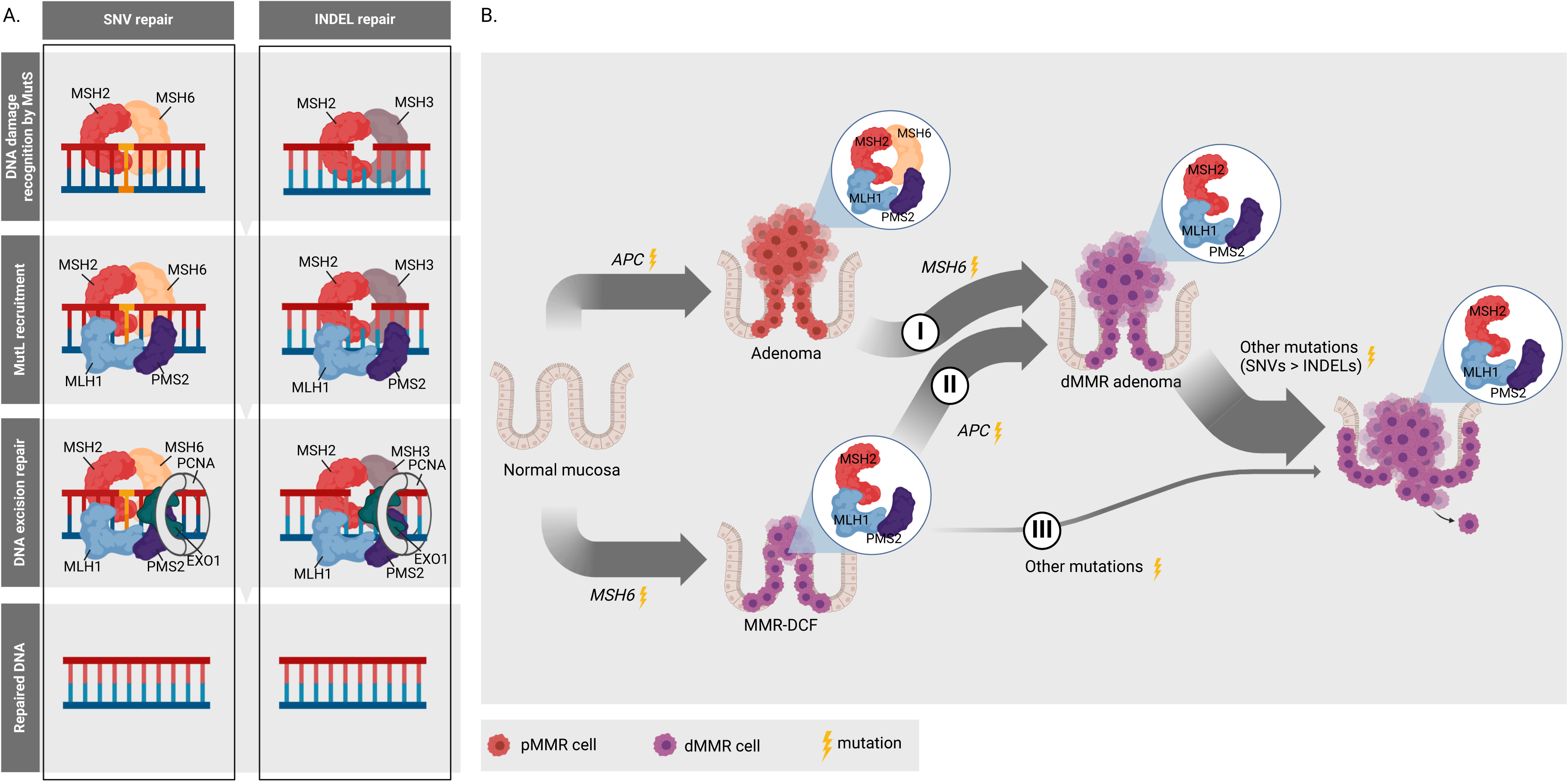
Function of MSH6 in relation to the development of *MSH6*-mutated CRCs. **A.** The repair of SNVs and INDELs by MMR is depicted. Briefly, MSH2 dimerizes with MSH6 (hMutSɑ) or with MSH3 (hMutSB) to scan the DNA for SNVs or INDELs, respectively. Once a SNV/INDEL is detected, hMutLɑ or hMutLɣ is recruited, consisting of MLH1 and PMS2 or MLH1 and MLH3, respectively. Next, hMutS-hMutL will trigger the degradation of the strand containing the DNA defect and will finally orchestrate the synthesis of a new strand.^9, 37–40^ In contrast to the other MMR proteins, MSH6 is thought to be redundant for the repair of INDELs, which mainly depends on the hMutSB.^41^ Adapted from Curtius et al.^40^ ^7, 9^ Created with BioRender.com. **B.** The development of LS CRCs through three molecular pathways, as proposed and adapted from Ahadova et al.^6^, is illustrated in the context of *MSH6*-mutated CRC development. The majority of *MSH6*-mutated CRCs is thought to originate from MMR-d adenomas, which in turn arise from MMR-proficient adenomas (I. adenoma – carcinoma pathway) or from MMR-DCF (II. MMR-DCF adenoma – carcinoma pathway).^9^ Our current findings indicate that the progression from MMR-d adenoma to carcinoma in *path_MSH6* carriers is likely driven primarily by SNVs rather than INDELs. This is based on the relatively lower abundance of cMS mutations in *MSH6*-mutated CRCs compared to other LS CRCs. Direct progression from MMR-DCF (III. MMR-DCF carcinoma pathway), excluding an intervening adenoma stage, has mainly been associated with *MLH1* carcinogenesis and is expected to play a minor role in *MSH6* carcinogenesis.^7, 9^ Created with BioRender.com. *cMS, coding microsatellite; CRC, colorectal cancer; FSP, frameshift peptide; HLA, human leukocyte antigen; INDEL, insertion/deletion variant; mAR, mutant allele ratio; (d)MMR, mismatch repair (deficiency/deficient); MMR-DCF, mismatch repair deficient crypt focus; SNV, single nucleotide variant*.

Secondly, our findings prompt the hypothesis that *MSH6*-mutated CRCs might elicit altered immune responses compared to CRCs from other MMR groups, considering the anticipated fewer FSPs derived from cMS mutations. Comprehensive immune profiling of LS CRCs stratified by MMR group would be essential to test this hypothesis and may shed further light on the influence of the immune system on the different pathways of colorectal carcinogenesis in *path_MMR* carriers.

In theory, altered immune responses directed towards *MSH6*-mutated CRCs could be reflected in the sensitivity to immunotherapies. However, as clinical studies to date have not described such differences, this aspect should be addressed in future larger trials assessing possible markers of therapy resistance. One approach that might be particularly affected by the lower mutation frequencies and mARs in *MSH6*-mutated CRCs involves FSP neoantigen-based vaccines. These vaccines target tumor cells with cancer-driving cMS mutations shared amongst LS-associated CRCs, and have shown promising potential in preclinical and clinical studies.^17, 18^ Our findings of lower cMS mutation frequencies and mARs in *MSH6*-mutated CRCs compared to CRCs from other MMR groups underline the need to account for *MSH6* cases in selecting FSP neoantigens for vaccines to obtain optimal coverage.

Potential target candidates that might be effective for most *path_MSH6* carriers include *MARCKS, ASTE1, TAF1B* and *BANP* mutations, which are shared among 59%, 68%, 68% and 70% of the *MSH6*-mutated CRCs, respectively. *BANP* mutations are of particular interest due to their relatively high mutation frequencies and mARs across all MMR groups, highlighting the point of shared cMS mutations among all LS carriers. Subcutaneous vaccination of FSP neoantigens originating from mutated *ASTE1* and *TAF1B* genes already underwent evaluation in a phase I/IIa clinical trial.^18^ The results demonstrated humoral and cellular immune responses, with no severe adverse events. Other cMS mutations which were frequently shared across multiple MMR groups from our cohort and, therefore, could potentially be promising vaccine candidates include *SLC22A9, PTHLH* and *TGFBR2*.

The observed variation in cMS mutation patterns between tumor regions and between tumors and adjacent adenomatous tissues shows that there is also within-patient heterogeneity. The inclusion of more adenomatous tissue and other pre-cancerous lesions (e.g. MMR-DCF) in cMS mutation analysis would be necessary to determine whether some cMS mutations may only be associated with premalignant lesions and others may drive the progression from pre-cancerous lesions to cancer. A potential candidate for the latter involves *ZNF294*, which encodes for Listerin E3 Ubiquitin Protein Ligase 1 (LTN1) and was mutated in three CRCs but not in the adjacent adenomatous tissues. This information could be relevant for identifying novel vaccine targets and determining appropriate administrative timeframes for prophylactic vaccination strategies.

Another factor that might be considered in the development of prophylactic vaccination strategies, involves the potential counterselection of cell clones with highly immunogenic FSPs, as implicated by the inverse correlations between the cMS mutation frequencies and the predicted immunogenicity of the resulting FSPs. This correlation has been described earlier^23, 24^, and might suggest that the selection of vaccine targets should not solely rely on the immunogenicity of the FSPs. To gain a better understanding of the tumor-immune cell interplay in LS CRCs, fundamental studies exploring the HLA profile and corresponding presentation of cMS mutation-derived FSPs in greater detail, such as considering the entire set of HLA alleles in each person^35^, as well as the *B2M* status and HLA class I expression profiles, would be necessary.

Our findings may have significance not only for hereditary CRC but also for sporadic cases, where MSI testing and IHC for the MMR proteins are increasingly used as predictive biomarkers for immunotherapy response.^36^ Our results, which indicate that the cMS mutation pattern of tumors with MSH6 defects differs from other MMR-d CRCs, even when all tumors are classified as MSI, may suggest that biomarkers based solely on the presence or absence of MSI lack relevant information. This suggests that cMS mutation patterns and/or tumor mutational burden may be better predictors, especially for MMR-d cases involving MSH6 defects. Here, a better understanding of the immune responses in *path_MSH6* carriers could deliver further information on the potential immune therapy-predictive value of the cMS mutational load. Future studies that compare MSI/IHC data with cMS mutation patterns in sporadic CRCs and correlate these patterns with immunotherapy outcomes (stratified by MMR gene defects) are necessary to explore this aspect.

In conclusion, we observed considerably lower cMS mutation frequencies and mARs in *MSH6*-mutated CRCs than in CRCs from other MMR groups, including sporadic MSI CRCs. These findings suggest that *MSH6*-mutated CRCs exhibit a lower degree of MSI, potentially contributing to the later age of onset observed in these cases. At the same time, it may imply an altered immune response compared to what is typically observed in MSI CRCs, a factor that should be thoroughly evaluated in future studies. If confirmed, these results would strengthen considering LS as multiple gene-specific syndromes, potentially leading to gene-specific screening, prevention and treatment guidelines.

## Acknowledgements

The authors sincerely thank all patients and their families for their participation in this study. We gratefully acknowledge the PALGA-Group Collaborators/participating pathology centers for providing patient samples. The excellent technical assistance by Ricarda Mehr, Vera Fuchs, Jonathan Dörre, Nina Nelius, Marieke E. IJsselsteijn and Manon van der Ploeg are gratefully acknowledged. The authors thank Noel F.C.C. de Miranda for reviewing this work.

## Abbreviations

B2M: B2-microglobulin

cMS: coding microsatellite

CRC: colorectal cancer

FSP: frameshift peptide

GELS: general epitope likelihood score

HLA: human leukocyte antigen

ICB: immune checkpoint blockade

INDEL: insertion-deletion mutation

IQR: interquartile range

LS: Lynch syndrome

*MLH1*-PM: *MLH1* promotor hypermethylation

MMR(-d): mismatch repair(-deficiency/deficient)

MSI: microsatellite instability

MSS: microsatellite stable

OLL: overall ligand likelihood

*path_MMR,* (likely): pathogenic variant of one of the mismatch repair genes

*path_MLH1,* (likely): pathogenic variant of one of the *MLH1* gene alleles

*path_MSH2,* (likely): pathogenic variant of one of the *MSH2* gene alleles

*path_MSH6,* (likely): pathogenic variant of one of the *MSH6* gene alleles

*path_PMS2,* (likely): pathogenic variant of one of the *PMS2* gene alleles

SD: standard deviation

SNV: single nucleotide variant

---

**Supplementary Figure 1. Schematic representation of *HLA-A*\*02:01 typing after sequencing.** Since the reverse primer was used for sequencing, the sequence of the reverse strand was obtained from the electropherogram and converted into the sense strand. The nucleotide at position 78 indicated whether an (A) *HLA-A*\*02:01 (T) or (B) non-*HLA-A*\*02:01 (C) allele was present. In case of an overlapping signal of T and C at position 78, the sample was considered to be heterozygous for the *HLA-A*\*02:01 allele.

**Supplementary Figure 2. Age at CRC diagnosis and tumor diameter. A.** Age at CRC diagnosis [years] of *path_MSH6* carriers exhibited a bimodal distribution and was significantly higher (65.0 years, IQR 6.8-71.0) as compared to the age at CRC diagnosis of *path_MLH1* (48.0, IQR 41.0-60.3; *P*=.046), *path_MSH2* (51.0, IQR 38.5-58.8; *P*=.013) and *path_PMS2* (46.0, IQR 41.0-57.8; *P*=.005) carriers. **B.** Tumor diameter [cm] did not statistically differ between *MSH6*-mutated CRCs (5.5, IQR 3.3-7.0) and *MLH1* (3.8, IQR 2.0-5.0; *P*=1.0), *MLH1-*PM (4.9, IQR 2.6-7.0; *P*=1.0), *MSH2* (2.5, IQR 2.0-6.0; *P*=1.0) and *PMS2* (5.0, IQR 3.5-6.5; *P*=1.0) CRCs. Statistical differences between *MSH6*-mutated CRCs versus *MLH1-*, *MSH2-, PMS2*-mutated CRCs or *MLH1-*PM CRCs were evaluated using the Kruskal-Wallis test. Raw *P-*values were adjusted for the number of comparisons and outcomes using the Benjamini & Hochberg correction. *P-*values are two-tailed and considered statistically significant when *P*<.05. Only significant *P*-values are shown. Level of significance: 0.1234 (ns), 0.0332 (*), 0.0021 (**), 0.0002 (***), <0.0001 (****). *CRC, colorectal cancer; IQR, interquartile range; MLH1-PM, MLH1 promotor hypermethylation; path_MLH1, (likely) pathogenic variant of one of the MLH1 gene alleles; path_MSH2, (likely) pathogenic variant of one of the MSH2 gene alleles; path_MSH6, (likely) pathogenic variant of one of the MSH6 gene alleles; path_PMS2, (likely) pathogenic variant of one of the PMS2 gene alleles*.

**Supplementary Figure 3. cMS mutation patterns including data from Bajwa-Ten Broeke et al. (2021).** Heatmap depicting unsupervised hierarchic clustering of the mARs of the 20 analyzed cMS. All MSI CRCs from the current cohort were considered, as well as all MSI CRCs described in Bajwa-Ten Broeke et al.^22^ Annotation for the original cohort is included. *cMS, coding microsatellite; CRC, colorectal cancer; mAR, mutant allele ratio; MLH1-PM, MLH1 promotor hypermethylation; MMR, mismatch repair; MSI, microsatellite instability*.

**Supplementary Figure 4. Association between cMS mutation pattern and age at CRC diagnosis or tumor diameter. A.** No association between age at CRC diagnosis (years) and the number of mutated cMS per CRC was observed (*P*=.287; Pearson’s r 0.081, 95% CI −0.068 – 0.22). Data from the current cohort and from Bajwa-Ten Broeke et al.^22^ were included. **B.** No association between age at CRC diagnosis (years) and the mean mARs of all cMS per CRC was observed (*P*=.893; Pearson’s r 0.010, 95% CI −0.140 – 0.160). Data from the current cohort and from Bajwa-Ten Broeke et al. (2021) were included. **C.** No association between tumor diameter [cm] and the number of mutated cMS per CRC was observed (*P*=.741; Pearson’s r −0.033, 95% CI −0.224 – 0.161). Tumor diameters were only available for samples from the current cohort. **D.** No association between tumor diameter [cm] and the mean mARs of all cMS per CRC was observed (*P*=.849; Pearson’s r 0.019, 95% CI −0.174 – 0.211). Tumor diameters were only available for samples from the current cohort. *cMS, coding microsatellite; CRC, colorectal cancer; mAR, mutant allele ratio; MLH1-PM, MLH1 promotor hypermethylation; MMR, mismatch repair*.

**Supplementary Figure 5. Association between predicted immunogenicity of cMS mutation-derived FSPs and respective cMS mutation frequency.** Correlation between the number of predicted epitopes in M1 cMS mutation-derived FSPs (GELS) and the frequency of the respective M1 cMS mutations, categorized by binding affinity, *B2M* mutation status and *HLA-A*\*02:01 status. The GELS, adapted from Ballhausen et al.^23^, are based on MHC ligand prediction and the prevalence of the respective HLA allele in European Caucasians, with a conservative estimate of predicted HLA-binding probability of *p*_binding_ = 50%. *B2M, B2-microglobulin; cMS, coding microsatellite; FSP, frameshift peptide; GELS, general epitope likelihood score*.

