## Supplementary figures and images for "Lower degree of microsatellite instability in colorectal carcinomas from *MSH6*-associated Lynch syndrome patients"

### Supplementary Figure 1

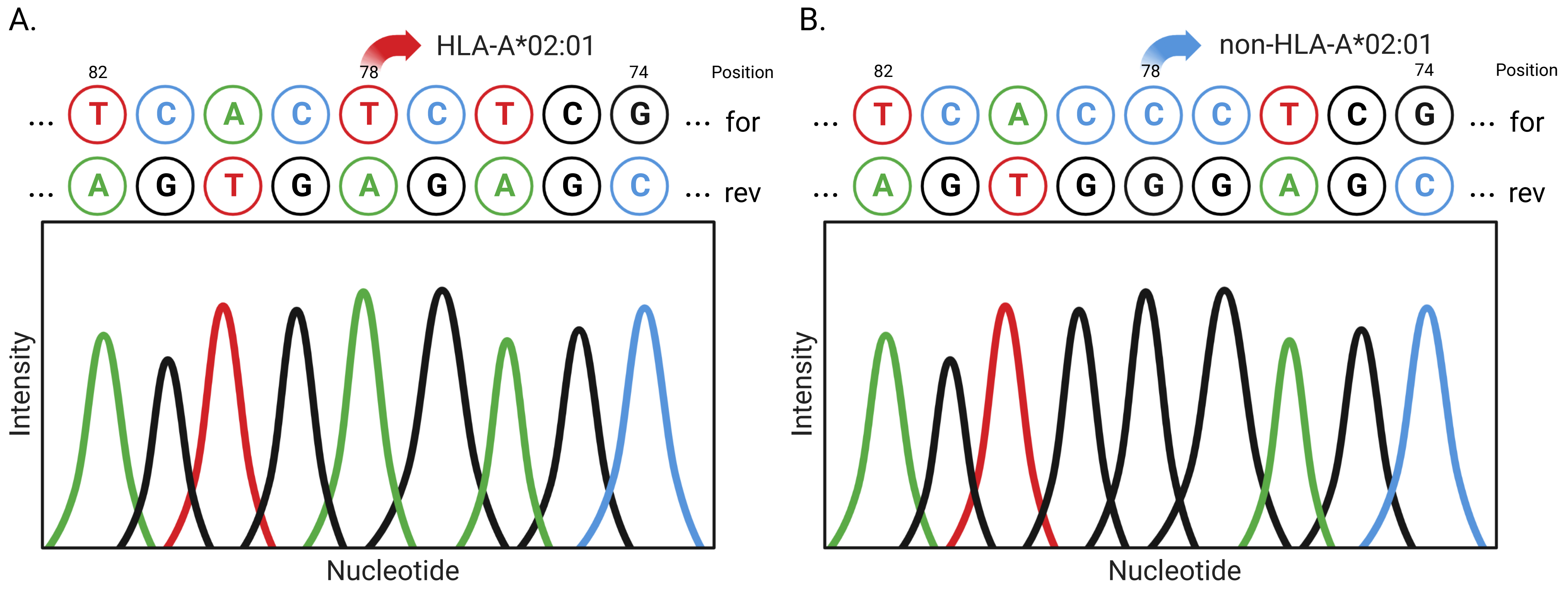

### Supplementary Figure 2

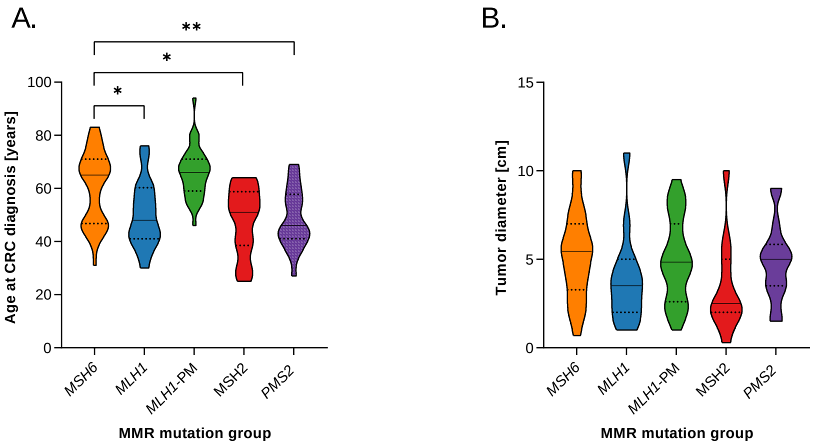

### Supplementary Figure 3

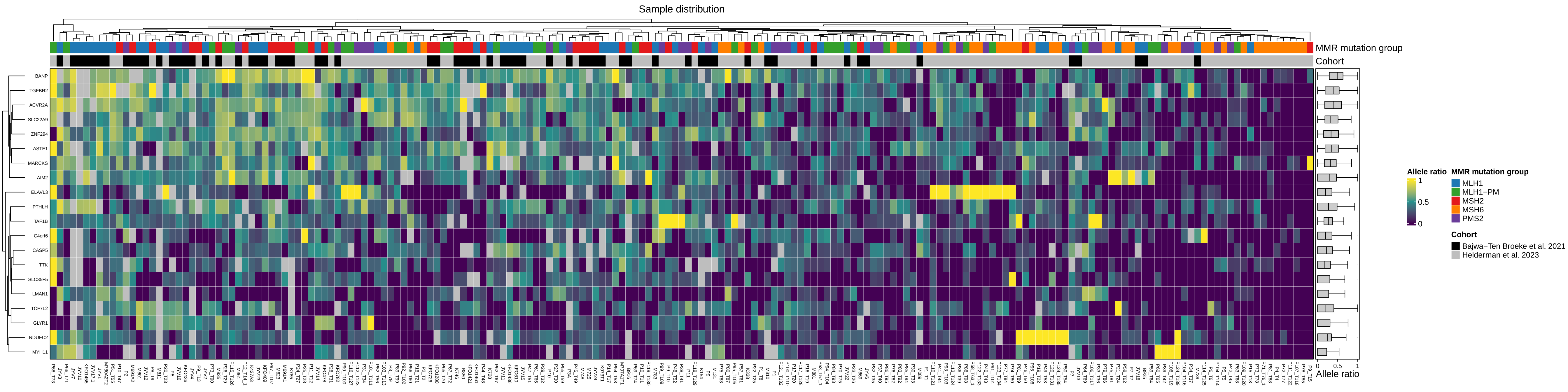

### Supplementary Figure 4

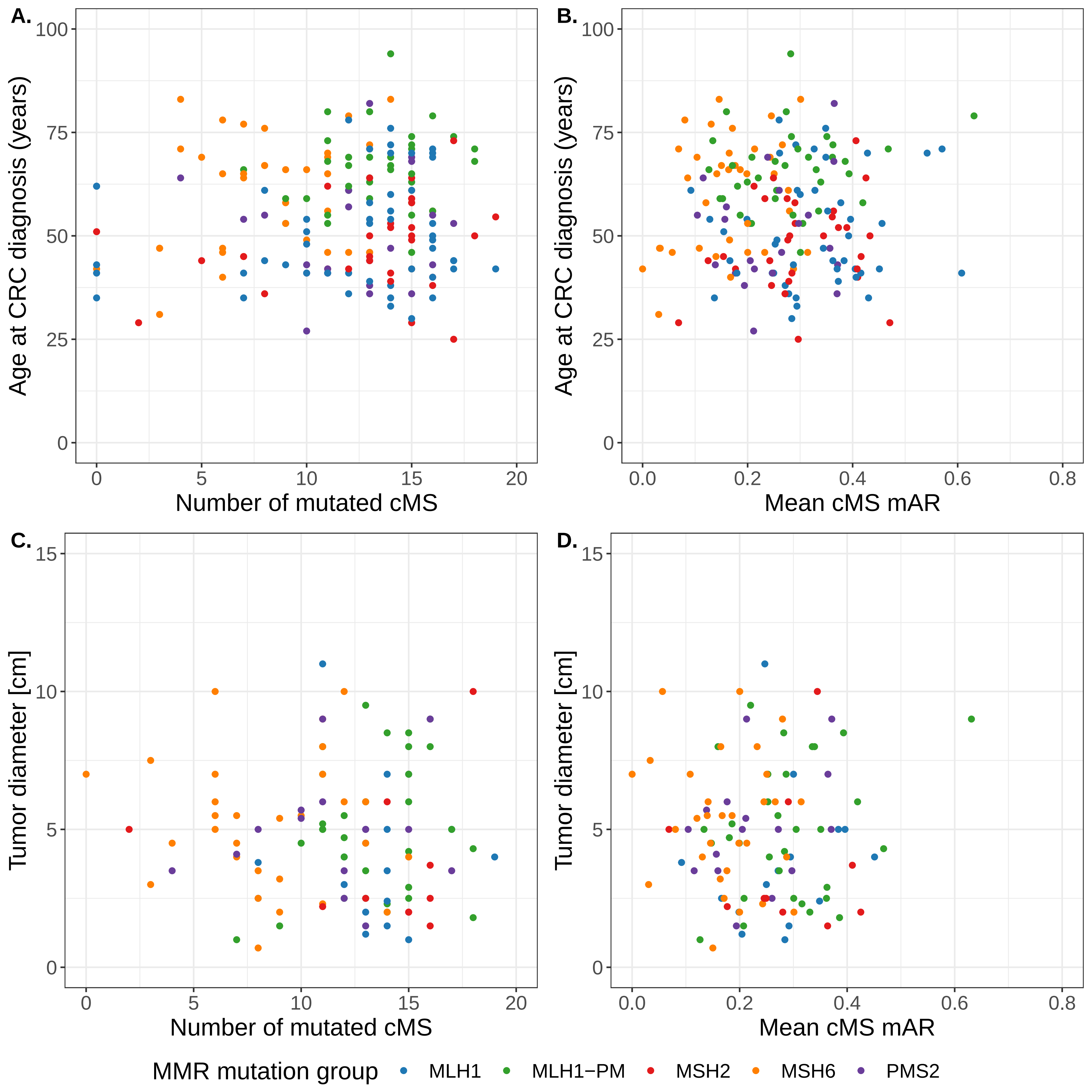

### Supplementary Figure 5

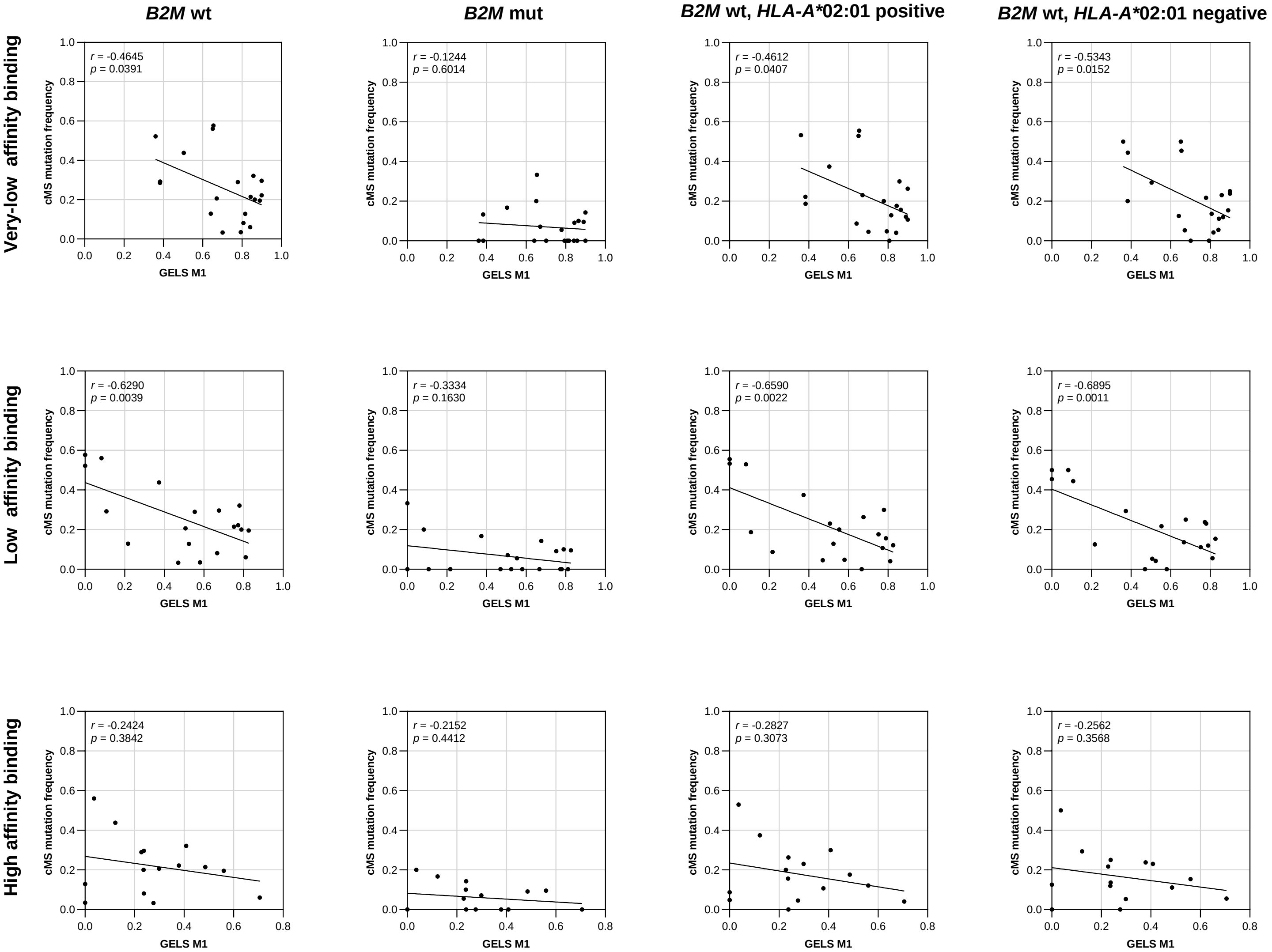
