## Supplementary Tables for "Lower degree of microsatellite instability in colorectal carcinomas from *MSH6*-associated Lynch syndrome patients"

### Supplementary Table 1. Materials and providers

| Device | Provider |
|---|---|
| 3500 XL Genetic Analyzer | Thermo Fisher Scientific, Waltham, MA, USA |
| ChemiDoc MP Imaging System | Bio-Rad Laboratories, Hercules, CA, USA |
| Consort EV231 Power Supply | Sigma-Aldrich, St. Louis, MO, USA |
| E1-ClipTip Electronic Multichannel Pipettes (1-30 µl, 2-125 µl) | Thermo Fisher Scientific, Waltham, MA, USA |
| Eppendorf Centrifuge 5810 R | Sigma-Aldrich, St. Louis, MO, USA |
| Fresco™ 21 Microcentrifuge | Thermo Fisher Scientific, Waltham, MA, USA |
| GeneTouch Thermal Cycler | Biozym Scientific, Hess. Oldendorf, Germany |
| Laminar Flow Hood, Blue Series | Kojair, Mänttä-Vilppula, Finland |
| Leica MZ6 Stereomicroscope | Leica Microsystems, Wetzlar, Germany |
| LifeTouch Thermal Cycler | Biozym Scientific, Hess. Oldendorf, Germany |
| Mettler Toledo PM4800 Precision Scale | Columbus, OH, USA |
| Nanodrop One | Thermo Fisher Scientific, Waltham, MA, USA |
| NanoZoomer S210 | Hamamatsu Photonics K.K.; Hamamatsu City, Japan |
| neoLab 3-1810 Mini-Centrifuge | neoLab Migge, Heidelberg, Germany |
| RS-OS 5 shaker | Phoenix Instrument, Garbsen, Germany |
| Savant SpeedVac | Thermo Fisher Scientific, Waltham, MA, USA |
| Single Channel Pipettes peqPETTE, (1000 µl, 200 µl, 20 µl) | PEQLAB Biotechnologie, Erlangen, Germany |
| Single Channel Pipettes Finnpipette F1 (1000 µl, 100 µl, 10 µl, 2 µl) | Thermo Fisher Scientific, Waltham, MA, USA |
| Sub-Cell GT Gel Electrophoresis Chamber | Bio-Rad Laboratories, Hercules, CA, USA |
| VF2 Vortex | IKA, Staufen im Breisgau, Germany |
| Heidolph Reax top vortex mixer | Heidolph Instruments, Schwabach, Germany |
| **Chemical/reagents** | **Provider** |
| 10 x PCR Rxn Buffer (-$MgCl_2$) | Thermo Fisher Scientific, Waltham, MA, USA |
| 100 mM dNTP Set | Thermo Fisher Scientific, Waltham, MA, USA |
| 50 mM $MgCl_2$ | Thermo Fisher Scientific, Waltham, MA, USA |
| Water, DNase, RNase-FREE | MP Biomedicals, Santa Ana, CA, USA |
| Platinum Taq-Polymerase | Thermo Fisher Scientific, Waltham, MA, USA |
| Taq-Polymerase | Thermo Fisher Scientific, Waltham, MA, USA |
| LE Agarose | Biozym Scientific, Hess. Oldendorf, Germany |
| 100 bp DNA Ladder | ThermoFisher Scientific, Waltham, USA |
| Xylene | ThermoFisher Scientific, Waltham, USA |
| Ethanol 100 % | Zentrallager Heidelberg University, Heidelberg, Germany |
| Ethanol 96 % | Zentrallager Heidelberg University, Heidelberg, Germany |
| Methanol | Honeywell, Charlotte, USA |
| Hydrogen peroxide (30 %) | Carl Roth, Karlsruhe, Germany |
| PBS (Gibco) | ThermoFisher Scientific, Waltham, USA |
| Tween 20 | Carl Roth, Karlsruhe, Germany |
| Normal Horse Serum (Vector) | BIOZOL Diagnostica Vertrieb, Eching, Germany |
| Solution A + B (Vectastain) | Vector Laboratories, Newark, CA, USA |
| DAB + Chromogen | Agilent Technologies, Santa Clara, CA, USA |
| DAB + Substrate Buffer | Agilent Technologies, Santa Clara, CA, USA |
| Eosin, yellowish | Waldeck, Münster, Germany |
| Mayer’s Hematoxylin | AppliChem, Darmstadt, Germany |
| EMSURE® ACS, ISO, Reag. Ph Eur Methanol for analysis | Merck Millipore, Burlington, MA, USA |
| **Consumable** | **Provider** |
| 96 Well PCR Platte, transparent | Biozym Scientific, Hess. Oldendorf, Germany |
| 96 Well PCR Platte, transparent, ABI Sequenzer 3100/30 | Biozym Scientific, Hess. Oldendorf, Germany |
| Aquatex | Merck, Darmstadt, Germany |
| BD Microlance Cannula | Becton Dickinson, Franklin Lakes, NJ, USA |
| Coverslips (multiple sizes) | Paul Marienfeld, Lauda Königshofen, Germany |
| Falcon tube (50 ml) | Corning Inc., Corning, NY, USA |
| Filter Tips (1000 µl, 200 µl) | Corning Inc., Corning, NY, USA |
| Filter Tips (10 µl) | Axygen |
| GeneScan 500 Rox Size Standard | ThermoFisher Scientific, Waltham, USA |
| HiDi Formamide | ThermoFisher Scientific, Waltham, USA |
| ImmEdge Hydrophobic Barrier PAP Pen | Vector Laboratories, Newark, CA, USA |
| Multichannel pipette tips (125 µl, 30 µl) | ThermoFisher Scientific, Waltham, USA |
| Precision wipes | Kimberly-Clark, Irving, TX, USA |
| primaSeal-qPCR | Steinbrenner Laborsysteme, Wiesenbach, Germany |
| POP-7 (960) | ThermoFisher Scientific, Waltham, USA |
| Reaction tubes (5 ml, 2 ml, 1.5 ml) | Eppendorf, Hamburg, Germany |
| Sapphire PCR 8-Tube Strips, 0.2 ml | Greiner Bio-One, Kremsmünster, Austria |
| **Kits** | **Provider** |
| BigDye Terminator v1.1 Cycle Sequencing Kit | ThermoFisher Scientific, Waltham, USA |

| | |
|---|---|
| QIAquick PCR Purification Kit REF: 28106 | Qiagen, Venlo, Netherlands |
| ReliaPrep FFPE gDNA Miniprep system REF: A2352 | Promega, Fitchburg, WI, USA |

| **cMS primers** | **Sequence (5' → 3')** | **Provider** |
|---|---|---|
| *ACVR2A* fwd | GTTGCCATTTGAGGAGGAAA | Thermo Scientific, Waltham, MA, USA |
| *ACVR2A* rev | CAGCATGTTTCTGCCAATAATC | Thermo Scientific, Waltham, MA, USA |
| *AIM2* fwd | TTCTCCATCCAGGTTATTAAGGC | Integrated DNA Technologies, Coralville, IA, USA |
| *AIM2* rev | TTAGACCAGTTGGCTTGAATTG | Integrated DNA Technologies, Coralville, IA, USA |
| *ASTE1* fwd | ATATGCCCCCGCTGAAATA | Microsynth, Balgach, Switzerland |
| *ASTE1* rev | TTGGTGTGTGCAGTGGTTCT | Microsynth, Balgach, Switzerland |
| *BANP* fwd | TTCTGTGGAAGCTCTGCCTT | Microsynth, Balgach, Switzerland |
| *BANP* rev | TCAAGTCGCATCAGATCCAG | Microsynth, Balgach, Switzerland |
| *C4orf6* fwd | CCAGAAGCAAATTCACAAGAC | Invitrogen, Waltham, MA, USA |
| *C4orf6* rev | TTTTGCGTGTTCCTTCCTTC | Invitrogen, Waltham, MA, USA |
| *CASP5* fwd | CAGAGTTATGTCTTAGGTGAAGG | Microsynth, Balgach, Switzerland |
| *CASP5* rev | ACCATGAAGAACATCTTTGCCCAG | Microsynth, Balgach, Switzerland |
| *ELAVL3* fwd | GATGCGACCTGTTATCTCCAG | Invitrogen, Waltham, MA, USA |
| *ELAVL3* rev | AGGTTGGTCTTGCTGTCGTC | Biomers.net, Ulm, Germany |
| *GLYR1* fwd | GCCTCCAGAAGCTGTGACTT | Invitrogen, Waltham, MA, USA |
| *GLYR1* rev | ATCACCAACATCCCGTCATT | Invitrogen, Waltham, MA, USA |
| *LMAN1* fwd | CACCCATGTCAGCTTTGCTA | Microsynth, Balgach, Switzerland |
| *LMAN1* rev | GGAGGAATTTGAGCACTTTCA | Microsynth, Balgach, Switzerland |
| *MARCKS* fwd | GACTTCTTCGCCCAAGGC | Invitrogen Waltham, MA, USA |
| *MARCKS* rev | GCCGCTCAGCTTGAAAGA | Biomers.net, Ulm, Germany |
| *MYH11* fwd | CGGGGATTCTCTCTCTGTTTC | Invitrogen, Waltham, MA, USA |
| *MYH11* rev | CTGAAGGCATGATACCTGGTG | Invitrogen, Waltham, MA, USA |
| *NDUFC2* fwd | TGAATTTCAGGTTTGCATCG | Sigma-Aldrich, St. Louis, MO, USA |
| *NDUFC2* rev | AACATTTCACGGTCCCTCAC | Thermo Electron Waltham, MA, USA |
| *PTHLH* fwd | TTTCACTTTCAGTACAGCACTTCTG | Thermo Scientific, Waltham, MA, USA |
| *PTHLH* rev | GAAGTAACAGGGGACTCTTAAATAATG | Biomers.net, Ulm, Germany |
| *SLC22A9* fwd | GCGCCTACAGTGCCTACTCT | Microsynth, Balgach, Switzerland |
| *SLC22A9* rev | GCATGTGGAGCATTTCACAC | Microsynth, Balgach, Switzerland |
| *SLC35F5* fwd | TGTGGGGAAACTTACTGCAA | Sigma-Aldrich, St. Louis, MO, USA |
| *SLC35F5* rev | TCAAGTTTCAAACATCATATGCAA | Integrated DNA Technologies, Coralville, IA, USA |
| *TAF1B* fwd | ACCCAAATAAAAGCCCTCAAC | Thermo Scientific Waltham, MA, USA |
| *TAF1B* rev | CTACTTAAAATTCCATTCCATGTCC | Integrated DNA Technologies, Coralville, IA, USA |
| *TCF7L2* fwd | GCCTCTATTCACAGATAACTC | Microsynth, Balgach, Switzerland |
| *TCF7L2* rev | GTTCACCTTGTATGTAGCGAA | Microsynth, Balgach, Switzerland |
| *TGFBR2* fwd | GCTGCTTCTCCAAAGTGCAT | Microsynth, Balgach, Switzerland |
| *TGFBR2* rev | CAGATCTCAGGTCCCACACC | Microsynth, Balgach, Switzerland |
| *TTK* fwd | TTCTTCATCCTCCAAGACTTTT | Microsynth, Balgach, Switzerland |
| *TTK* rev | GATTTCCACAGGGATTCAAGA | Invitrogen, Waltham, MA, USA |
| *ZNF294* fwd | AAGCCGAAGAGCTCATTGAA | Microsynth, Balgach, Switzerland |
| *ZNF294* rev | CAGTTGTTAATTCCCAGCCTTC | Microsynth, Balgach, Switzerland |
| ***B2M* primers** | **Sequence (5' → 3')** | **Provider** |
| B2M Exon 1 F | GGCATTCCTGAAGCTGACA | Life Technologies, Darmstadt, Germany |
| B2M Exon 1 R | AGAGCGGGAGAGGAAGGAC | Life Technologies, Darmstadt, Germany |
| B2M Exon 2 F | TGACACCAAGTTAGCCCCAA | Life Technologies, Darmstadt, Germany |
| B2M Exon 2 R | ACTCATACACAACTTTCAGCAGC | Life Technologies, Darmstadt, Germany |
| ***HLA-A**02:01 primers** | **Sequence (5' → 3')** | **Provider** |
| Vilabona/A2-2F | TCTCAGCCACTCCTCGTC | Integrated DNA Technologies, Coralville, IA, USA |
| G R | TGTCGAACCGCACGAACTG | Integrated DNA Technologies, Coralville, IA, USA |
| **Antibody [clone]** | **Dilution** | **Provider** |
| MLH1 [G168-15] Purified Mouse Anti-Human | 1:50 | BD Biosciences, Franklin Lakes, NJ, USA |
| MSH2 [FE11] (Ab-2) Mouse mAb | 1:200 | Calbiochem, San Diego, CA, USA |
| MSH6 [EP49] Rabbit Monoclonal Anti-Human | 1:200 | Epitomics, Franklin Lakes, CA, USA |
| PMS2 [EP51] RMab | 1:75 | Bio SB, Santa Barbara, CA, USA |
| Horse Anti-Mouse/Rabbit IgG (H+L), Biotinylated, R.T.U. | 1:1 | Vector Laboratories, Newark, CA, USA |
| **Software** | **Description** | **Provider** |

| | | |
|---|---|---|
| 3500 Series Data Collection Software 3 | Software for 3500 XL Genetic Analyzer | Applied Biosystems; Waltham, USA |
| Gene Marker v1.50 | Analysis of results from fragment length analysis | SoftGenetics; State College, USA |
| Image Lab v6.1.0 | Gel electrophoresis imaging | Bio-Rad Laboratories, Hercules, CA, USA |
| NDP.scan v3.2.15 | Scanning of IHC slides | Hamamatsu Photonics K.K.; Hamamatsu City, Japan |
| NDP.view v2.7.25 | Viewing of IHC slides | Hamamatsu Photonics K.K.; Hamamatsu City, Japan |
| R v4.1.3 | Statistical analysis and ReFrame algorithm | R Foundation for Statistical Computing, Vienna, Austria |
| RStudio v2022.02.3+492 | Statistical analysis and ReFrame algorithm | RStudio Team, Boston, MA, USA |
| IBM SPSS Statistics v29.0 | Statistical analysis | International Business Machines Corporation (IBM), NY, USA |
| Sequencing Analysis Software v7.1 | Analysis of sequencing results | Applied Biosystems; Waltham, USA |
| CaseViewer v2.4.0 | | 3DHISTECH Ltd., Budapest, Hungary |

### Supplementary Table 2. MMR/MSI status

| MMR mutation group | IHC MMR[a] | | | MSI[b,c] | | | BAT25 | | | BAT26 | | | CAT25 | | | BAT40 | | |
|---|---|---|---|---|---|---|---|---|---|---|---|---|---|---|---|---|---|---|
| | # interpretable data | # loss | % loss | # interpretable data | # instable | % instable | # interpretable data | # instable | % instable | # interpretable data | # instable | % instable | # interpretable data | # instable | % instable | # interpretable data | # instable | % instable |
| ***MSH6* (n=39)** | 32 | 29 | 90% | 39 | 38 | 97% | 38 | 34 | 89% | 37 | 34 | 92% | 26 | 24 | 92% | 32 | 31 | 97% |
| ***MLH1* (n=18)** | 18 | 18 | 100% | 18 | 18 | 100% | 16 | 16 | 100% | 16 | 16 | 100% | 15 | 15 | 100% | 14 | 14 | 100% |
| ***MLH1*-PM (n=35)** | 35 | 35 | 100% | 35 | 35 | 100% | 23 | 23 | 100% | 23 | 23 | 100% | 19 | 19 | 100% | 18 | 18 | 100% |
| ***MSH2* (n=16)** | 16 | 16 | 100% | 16 | 15 | 94% | 15 | 14 | 93% | 15 | 13 | 87% | 14 | 13 | 93% | 14 | 13 | 93% |
| ***PMS2* (n=22)** | 7 | 6 | 86% | 22 | 20 | 91% | 26 | 24 | 92% | 16 | 13 | 81% | 14 | 12 | 86% | 14 | 12 | 86% |

[a]MMR protein loss corresponding with the constitutional MMR variant.
[b]MSI status was extracted from pathology records (n=40) or was evaluated by (multiplex) PCR and fragment length analysis (BAT25, BAT26, CAT25, BAT40) in case not reported in pathology records (n=90). For 23 CRCs that were MSI according to the pathology records, specific marker data was not available.
[c]CRCs were considered MSI if 2 ≥ markers were instable.

*CRC, colorectal cancer; IHC, immunohistochemistry; MLH1-PM, MLH1 promotor hypermethylation; MMR, mismatch repair; MSI, microsatellite instability*

### Supplementary Table 3. cMS mutation frequencies per MMR mutation group including Bajwa-Ten Broeke et al. 2021 data

| cMS | MMR mutation groups | | | | | | | | | | *P* value[a] | | | | | | | | | |
|---|---|---|---|---|---|---|---|---|---|---|---|---|---|---|---|---|---|---|---|---|
| | *MSH6* (n=38) | | *MLH1* (n=59) | | *MLH1-PM* (n=35) | | *MSH2* (n=36) | | *PMS2* (n=32) | | Overall | | *MLH1vsMSH6* | | *MLH1-PMvsMSH6* | | *MSH2vsMSH6* | | *PMS2vsMSH6* | |
| | n | % | n | % | n | % | n | % | n | % | Raw | Adj. | Raw | Adj. | Raw | Adj. | Raw | Adj. | Raw | Adj. |
| ***ACVR2A*** | 17/38 | 45% | 45/51 | 88% | 29/34 | 85% | 28/33 | 85% | 26/30 | 87% | **<0.001** | **<0.001** | **<0.001** | **0.001** | **<0.001** | **0.028** | **<0.001** | **0.037** | **<0.001** | **0.030** |
| ***AIM2*** | 14/37 | 38% | 36/48 | 75% | 27/34 | 79% | 31/32 | 97% | 19/30 | 63% | **<0.001** | **<0.001** | **0.001** | **0.045** | **<0.001** | **0.032** | **<0.001** | **<0.001** | **0.038** | 1 |
| ***ASTE1*** | 26/38 | 68% | 42/48 | 88% | 25/32 | 78% | 27/32 | 84% | 23/31 | 74% | 0.226 | 1 | **0.031** | 1 | 0.363 | 1 | 0.121 | 1 | 0.599 | 1 |
| ***BANP*** | 26/38 | 68% | 42/46 | 91% | 34/35 | 97% | 29/30 | 97% | 27/31 | 87% | **0.001** | **0.017** | **0.008** | 0.628 | **0.001** | 0.108 | **0.003** | 0.262 | 0.067 | 1 |
| ***C4orf6*** | 16/37 | 43% | 29/48 | 60% | 21/35 | 60% | 15/32 | 47% | 20/28 | 71% | 0.145 | 1 | 0.116 | 1 | 0.155 | 1 | 0.762 | 1 | **0.024** | 1 |
| ***CASP5*** | 13/37 | 35% | 35/46 | 76% | 25/34 | 74% | 22/31 | 71% | 12/29 | 41% | **<0.001** | **0.003** | **<0.001** | **0.014** | **0.001** | 0.096 | **0.003** | 0.259 | 0.604 | 1 |
| ***ELAVL3*** | 14/37 | 38% | 27/47 | 57% | 20/35 | 57% | 20/31 | 65% | 19/29 | 66% | 0.137 | 1 | 0.074 | 1 | 0.101 | 1 | **0.028** | 1 | **0.026** | 1 |
| ***GLYR1*** | 7/38 | 18% | 29/51 | 57% | 10/35 | 29% | 17/32 | 53% | 21/30 | 70% | **<0.001** | **0.001** | **<0.001** | **0.021** | 0.305 | 1 | **0.002** | 0.185 | **<0.001** | **0.001** |
| ***LMAN1*** | 6/38 | 16% | 27/52 | 52% | 17/35 | 49% | 17/31 | 55% | 11/30 | 37% | **0.003** | 0.063 | **<0.001** | **0.035** | **0.003** | 0.208 | **0.001** | **0.050** | **0.048** | 1 |
| ***MARCKS*** | 22/37 | 59% | 38/45 | 84% | 28/35 | 80% | 26/30 | 87% | 21/30 | 70% | **0.038** | 0.759 | **0.011** | 0.884 | 0.059 | 1 | **0.014** | 1 | 0.371 | 1 |
| ***MYH11*** | 6/38 | 16% | 19/48 | 40% | 15/35 | 43% | 10/33 | 30% | 9/22 | 41% | 0.084 | 1 | **0.016** | 1 | **0.011** | 0.856 | 0.144 | 1 | **0.030** | 1 |
| ***NDUFC2*** | 17/37 | 46% | 40/47 | 85% | 22/35 | 63% | 23/30 | 77% | 21/32 | 66% | **0.003** | 0.057 | **<0.001** | **0.011** | 0.150 | 1 | **0.011** | 0.864 | 0.101 | 1 |
| ***PTHLH*** | 18/38 | 47% | 38/50 | 76% | 30/35 | 86% | 21/31 | 68% | 27/31 | 87% | **0.001** | **0.015** | **0.006** | 0.454 | **0.001** | **0.045** | 0.089 | 1 | **0.001** | **0.045** |
| ***SLC22A9*** | 19/38 | 50% | 36/46 | 78% | 30/35 | 86% | 29/31 | 94% | 25/31 | 81% | **<0.001** | **0.004** | **0.007** | 0.536 | **0.001** | 0.094 | **<0.001** | **0.007** | **0.008** | 0.674 |
| ***SLC35F5*** | 8/38 | 21% | 25/48 | 52% | 14/35 | 40% | 22/32 | 69% | 18/31 | 58% | **0.001** | **0.015** | **0.003** | 0.264 | 0.078 | 1 | **<0.001** | **0.005** | **0.002** | 0.128 |
| ***TAF1B*** | 26/37 | 70% | 39/50 | 78% | 26/33 | 79% | 25/34 | 74% | 23/30 | 77% | 0.909 | 1 | 0.412 | 1 | 0.416 | 1 | 0.760 | 1 | 0.557 | 1 |
| ***TCF7L2*** | 11/38 | 29% | 29/43 | 67% | 11/35 | 31% | 22/32 | 69% | 19/27 | 70% | **<0.001** | **0.001** | **0.001** | **0.044** | 0.817 | 1 | **0.001** | 0.071 | **0.001** | 0.077 |
| ***TGFBR2*** | 16/38 | 42% | 43/45 | 96% | 31/34 | 91% | 28/30 | 93% | 19/26 | 73% | **<0.001** | **<0.001** | **<0.001** | **<0.001** | **<0.001** | **0.001** | **<0.001** | **0.001** | **0.015** | 1 |
| ***TTK*** | 15/38 | 39% | 27/47 | 57% | 21/35 | 60% | 22/31 | 71% | 9/22 | 41% | 0.059 | 1 | 0.099 | 1 | 0.080 | 1 | **0.009** | 0.726 | 0.913 | 1 |
| ***ZNF294*** | 19/38 | 50% | 47/52 | 90% | 28/35 | 80% | 28/33 | 85% | 20/31 | 65% | **<0.001** | **0.002** | **<0.001** | **0.002** | **0.007** | 0.600 | **0.002** | 0.157 | 0.226 | 1 |

[a]cMS mutation frequencies were compared between the MMR mutation groups using Chi-square tests and raw *P* values were subsequently adjusted for the number of comparisons and outcomes under investigation using Benjamini & Hochberg correction for multiple testing. Raw *P* values from the overall comparison were corrected for 20 tests (one comparison and 20 outcomes), whereas the raw *P* values from the pairwise comparisons were corrected for 80 tests (four comparisons and 20 outcomes). *P* values are two-tailed and considered statistically significant when *P* < .05. Significant *P* values are marked bold.

*cMS, coding microsatellite; MLH1-PM; MLH1 promotor hypermethylation; MMR, mismatch repair*

### Supplementary Table 4. cMS mARs per MMR mutation group including data from Bajwa-Ten Broeke et al. 2021

| cMS | mAR per MMR group | | | | | *P* value[a] | | | | | | | | | |
|---|---|---|---|---|---|---|---|---|---|---|---|---|---|---|---|
| | *MSH6* (n=38) | *MLH1* (n=59) | *MLH1-PM* (n=35) | *MSH2* (n=36) | *PMS2* (n=32) | Overall | | *MLH1*vsMSH6 | | *MLH1-PMvsMSH6* | | *MSH2vsMSH6* | | *PMS2*vsMSH6 | |
| | | | | | | Raw | Adj. | Raw | Adj. | Raw | Adj. | Raw | Adj. | Raw | Adj. |
| ***ACVR2A*** | 0.212 | 0.464 | 0.419 | 0.439 | 0.402 | **<0.001** | **0.002** | **<0.001** | **<0.001** | **0.002** | **0.047** | **0.001** | **0.017** | **0.009** | 0.171 |
| ***AIM2*** | 0.154 | 0.339 | 0.337 | 0.434 | 0.271 | **<0.001** | **0.003** | **0.003** | 0.068 | **0.009** | 0.180 | **<0.001** | **0.001** | 0.181 | 1 |
| ***ASTE1*** | 0.239 | 0.42 | 0.39 | 0.38 | 0.284 | **0.002** | **0.048** | **0.002** | **0.031** | **0.024** | 0.489 | **0.040** | 0.802 | 0.838 | 1 |
| ***BANP*** | 0.264 | 0.527 | 0.545 | 0.557 | 0.391 | **<0.001** | **<0.001** | **<0.001** | **<0.001** | **<0.001** | **<0.001** | **<0.001** | **<0.001** | 0.089 | 1 |
| ***C4orf6*** | 0.208 | 0.237 | 0.225 | 0.157 | 0.286 | **0.047** | 0.939 | 0.134 | 1 | 0.287 | 1 | 0.990 | 1 | **0.030** | 0.599 |
| ***CASP5*** | 0.116 | 0.296 | 0.283 | 0.245 | 0.157 | **<0.001** | **0.008** | **<0.001** | **0.009** | **0.003** | 0.066 | **0.040** | 0.794 | 0.840 | 1 |
| ***ELAVL3*** | 0.248 | 0.212 | 0.298 | 0.207 | 0.323 | 0.381 | 1 | 1 | 1 | 0.666 | 1 | 0.999 | 1 | 0.473 | 1 |
| ***GLYR1*** | 0.058 | 0.23 | 0.104 | 0.201 | 0.297 | **<0.001** | **0.001** | **0.001** | **0.026** | 0.788 | 1 | **0.026** | 0.511 | **<0.001** | **0.001** |
| ***LMAN1*** | 0.057 | 0.184 | 0.192 | 0.169 | 0.133 | **0.025** | 0.509 | **0.014** | 0.282 | **0.018** | 0.367 | 0.078 | 1 | 0.355 | 1 |
| ***MARCKS*** | 0.176 | 0.374 | 0.353 | 0.425 | 0.262 | **<0.001** | **0.001** | **0.001** | **0.011** | **0.005** | 0.095 | **<0.001** | **0.001** | 0.359 | 1 |
| ***MYH11*** | 0.134 | 0.157 | 0.162 | 0.094 | 0.162 | 0.500 | 1 | 0.555 | 1 | 0.559 | 1 | 1 | 1 | 0.668 | 1 |
| ***NDUFC2*** | 0.206 | 0.347 | 0.227 | 0.238 | 0.185 | **0.022** | 0.439 | **0.028** | 0.559 | 0.988 | 1 | 0.953 | 1 | 0.989 | 1 |
| ***PTHLH*** | 0.171 | 0.355 | 0.35 | 0.269 | 0.311 | **0.002** | **0.049** | **0.001** | **0.019** | **0.004** | 0.077 | 0.234 | 1 | **0.042** | 0.848 |
| ***SLC22A9*** | 0.226 | 0.342 | 0.373 | 0.422 | 0.33 | **0.012** | 0.242 | 0.085 | 1 | **0.031** | 0.618 | **0.003** | 0.058 | 0.215 | 1 |
| ***SLC35F5*** | 0.084 | 0.186 | 0.14 | 0.227 | 0.192 | **0.034** | 0.686 | 0.072 | 1 | 0.596 | 1 | **0.014** | 0.278 | 0.095 | 1 |
| ***TAF1B*** | 0.248 | 0.262 | 0.267 | 0.284 | 0.351 | 0.429 | 1 | 0.997 | 1 | 0.992 | 1 | 0.919 | 1 | 0.226 | 1 |
| ***TCF7L2*** | 0.126 | 0.318 | 0.092 | 0.274 | 0.279 | **<0.001** | **<0.001** | **<0.001** | **0.010** | 0.909 | 1 | **0.020** | 0.397 | **0.023** | 0.455 |
| ***TGFBR2*** | 0.177 | 0.441 | 0.376 | 0.514 | 0.375 | **<0.001** | **<0.001** | **<0.001** | **<0.001** | **0.002** | **0.032** | **<0.001** | **<0.001** | **0.004** | 0.081 |
| ***TTK*** | 0.094 | 0.186 | 0.189 | 0.255 | 0.144 | **0.009** | 0.189 | 0.084 | 1 | 0.102 | 1 | **0.002** | **0.036** | 0.730 | 1 |
| ***ZNF294*** | 0.174 | 0.436 | 0.328 | 0.434 | 0.262 | **<0.001** | 0.086 | **<0.001** | **<0.001** | **0.016** | 0.319 | **<0.001** | **<0.001** | 0.322 | 1 |

[a]cMS mARs were compared by one-way ANOVA with Dunnett's test for post-hoc analysis. For the pairwise comparisons, raw *P* values were additionally corrected for the number of comparisons (n=4) using Benjamini & Hochberg correction for multiple testing. *P* values are two-tailed and considered statistically significant when *P* < .05. Significant *P* values are marked bold.

*cMS, coding microsatellite; mAR, mutant allele ratio; MLH1-PM; MLH1 promotor hypermethylation; MMR, mismatch repair*

### Supplementary Table 5. cMS mARs of CRCs with multiple tumor regions or (adjacent) adenomatous tissue

| ID[a] | CRCs with multiple tumor regions | | | | | | | | | CRCs with (adjacent) adenomatous tissue | | | | | | | |
|---|---|---|---|---|---|---|---|---|---|---|---|---|---|---|---|---|---|
| | I | | II | | | III | | IV | | I | | II | | III | | IV | |
| | T1 | T2 | T1 | T2 | T3 | T1 | T2 | T1 | T2 | Ad | T | Ad | T | Ad | T | Ad | T |
| MMR | *MSH2* | | *MLH1* | | | *MSH6* | | *MSH6* | | *PMS2* | | *MLH1* | | *MLH1* | | *MSH6* | |
| ***ACVR2A*** | 0.66 | 0.25 | 0.4 | 0.61 | 0.17 | 0.4 | 0 | 0.4 | 0.2 | 0.61 | 0 | 0 | 0.38 | 0.43 | 0.49 | 1 | 0.68 |
| ***AIM2*** | 0.67 | 0.4 | 0.44 | 0.53 | 0.49 | 0 | 0 | 0 | 0 | 0.41 | 0 | 0.69 | | 0 | 0 | 0 | 0 |
| ***ASTE1*** | 0.65 | 0.22 | 0.2 | 0.27 | 0.5 | 0.37 | 0 | 0.21 | 0 | 1 | 0.62 | 0.2 | 0.53 | 0.46 | 0.58 | 0.28 | 0.29 |
| ***BANP*** | 0.76 | 0.26 | 0.62 | 0.8 | 0.62 | 0.52 | 0 | 0.27 | 0 | 0 | 0 | 1 | 0.6 | 0.47 | 0.69 | 0.48 | 0.69 |
| ***C4orf6*** | 0.44 | 0.67 | 0.45 | 0.28 | 0.42 | 0 | 0 | 0 | 0 | 0.35 | 0 | 0 | 0 | 0.66 | 0.64 | 0.28 | 0.42 |
| ***CASP5*** | 0.28 | 0 | 0.21 | 0.55 | 0 | 0.46 | 0 | 0 | 0 | 0 | 0 | 0 | 0 | 0 | 0 | 0 | 0.56 |
| ***ELAVL3*** | 0 | 0 | 0.25 | 0.4 | 0.27 | 1 | 0.18 | 0 | 1 | 1 | 0 | | 0 | 0 | 0.58 | 0 | 0 |
| ***GLYR1*** | 0.33 | 0.26 | 0.28 | 0.4 | 0.29 | 0 | 0 | 0 | 0 | 0 | 0 | 0.19 | 0 | 0 | 0 | 0 | 0 |
| ***LMAN1*** | 0 | 0 | 0 | 0 | 0 | 0 | 0 | 0 | 0 | 0 | 0 | 0 | 0.43 | 0 | 0 | 0 | 0 |
| ***MARCKS*** | 0.41 | 0 | 0.48 | 0.64 | 0.46 | 0 | 0 | 0.28 | 0 | 0.39 | 0 | 0 | 0.25 | 0.54 | 0.73 | 0 | 0 |
| ***MYH11*** | 0 | 0 | 0.34 | 0.69 | 0.43 | 0.2 | 0 | 0 | 0 | 0 | 1 | 0.17 | 0 | 0 | 0 | 0 | 0 |
| ***NDUFC2*** | 0.2 | 0 | 0 | 0 | 0.35 | 0 | 0 | 0.16 | 0.21 | 0.74 | 0 | 1 | 1 | 0.27 | 0.41 | 0 | 0 |
| ***PTHLH*** | | 0.46 | 0.26 | 0.33 | 0.58 | 0.45 | 0.23 | 0 | 0 | 0.25 | 0.29 | 0.5 | 0.34 | 0.29 | 0.33 | 0 | 0 |
| ***SLC22A9*** | 0.38 | 0.23 | 0.19 | 0.29 | 0.58 | 0.27 | 0 | 0 | 0.35 | 0.69 | 0 | 0.17 | 0.26 | 0.54 | 0.69 | 0 | 0 |
| ***SLC35F5*** | 0.27 | 0.15 | 0 | 0 | 0 | 0 | 0 | 0 | 0 | 0.19 | 0 | 0 | 0 | 0.27 | 0.37 | 0 | 0 |
| ***TAF1B*** | 0.34 | 0.16 | 0 | 0.32 | 0.28 | 0.51 | 0.19 | 0.24 | 0 | 0 | 0 | 0.59 | 0.2 | 0.24 | 0.28 | 0 | |
| ***TCF7L2*** | 0.45 | 0.28 | 0.27 | 0.39 | 0 | 0 | 0 | 0 | 0 | 0 | 0 | 0 | 0.25 | 0 | 0 | 0 | 0 |
| ***TGFBR2*** | 0.74 | 0 | 0.32 | 1 | 0.29 | 0 | 0.55 | 0 | 0 | 0 | 0.4 | 0.29 | 0.32 | 0.58 | 0.6 | 0 | 0 |
| ***TTK*** | 0.48 | 0.26 | 0.18 | 0 | 0 | 0.16 | 0 | 0.16 | 0 | 0.15 | 0 | 0 | 0 | 0.33 | 0.32 | 0 | 0 |
| ***ZNF294*** | 0.72 | 0.35 | 0.23 | 0.25 | 0.51 | 0.31 | 0.42 | 0 | 0.24 | 0 | 0 | 0 | 0.18 | 0 | 0.27 | 0 | 0.3 |

[a]IDs correspond with Supplementary Figure 5.

*Ad, adenoma; cMS, coding microsatellite; CRC, colorectal cancer; mAR, mutant allele ratio*

### Supplementary Table 6. Overall ligand likelihood and m1-frameshift mutation frequency of every cMS

| cMS | GELS[a] | | | $OLL_{HLA\text{-}A2*02:01}$-$OLL_{HLA\text{-}A2*02:01}$[b] | $m_{HLA\text{-}A2*02:01}$-$m_{HLA\text{-}A2*02:01}$ | |
|---|---|---|---|---|---|---|
| | **Very-low affinity binding** | **Low affinity binding** | **High affinity binding** | | **All *MSH6* CRCs** (n=38) | ***B2M* wildtype *MSH6* CRCs** (n=22) |
| ***ACVR2A*** | 0.36082841140750815 | 0.0 | | -0.00930 | 0.00132 | -0.1193033484 |
| ***AIM2*** | 0.3834326780087304 | 0.10785632 | | -0.08786 | -0.07311 | -0.0766366111 |
| ***ASTE1*** | 0.6509486752372561 | 0.08324785751978225 | 0.036604355 | 0.01540 | 0.06302 | 0.0114421718 |
| ***BANP*** | 0.6544460967564362 | 0.0 | | 0.07554 | -0.02160 | -0.1075557373 |
| ***C4orf6*** | 0.8993831291141262 | 0.6762411763203415 | 0.23754987500000002 | 0.40371 | -0.02884 | -0.0426199590 |
| ***CASP5*** | 0.8578973828005656 | 0.7792468958164972 | 0.40847629021001625 | 0.26267 | -0.04806 | -0.0006531228 |
| ***ELAVL3*** | 0.793398826152717 | 0.5801188652102128 | 0.0 | 0.68741 | 0.04585 | 0.0134666206 |
| ***GLYR1*** | 0.7013075395409308 | 0.4704780976578697 | 0.2764552571785448 | -0.26135 | 0.03856 | 0.0194379162 |
| ***LMAN1*** | 0.6411866806798175 | 0.21676192271909758 | 0.0 | -0.27023 | -0.03248 | -0.0860771624 |
| ***MARCKS*** | 0.5039459196624102 | 0.3736818049251361 | 0.12207182 | -0.03072 | -0.02113 | -0.0341017373 |
| ***MYH11*** | 0.8415748352142511 | 0.8111131593283832 | 0.7057856894606713 | 0.25922 | 0.03159 | 0.0365318033 |
| ***NDUFC2*** | 0.6711674731857592 | 0.507433183782016 | 0.2993246668066775 | -0.01050 | 0.09026 | 0.0544310105 |
| ***PTHLH*** | 0.38293081005492424 | | | -0.07540 | -0.03378 | -0.0743032885 |
| ***SLC22A9*** | 0.8999091401427837 | 0.7726600634108375 | 0.37883728809352807 | 0.32412 | -0.07786 | -0.1915897254 |
| ***SLC35F5*** | 0.8068003922354917 | 0.6673483331390249 | 0.23754987500000002 | -0.08966 | -0.11094 | -0.1601827135 |
| ***TAF1B*** | 0.7786799115713393 | 0.5534754620903691 | 0.22769949125 | -0.26728 | 0.01463 | -0.0665778584 |
| ***TCF7L2*** | 0.8158820363971048 | 0.5248877197163252 | | -0.27162 | -0.00733 | 0.0466167863 |
| ***TGFBR2*** | 0.8643826364016642 | 0.7894164498616978 | 0.2363025628228791 | 0.17419 | -0.04248 | -0.0676693133 |
| ***TTK*** | 0.8906920741790854 | 0.8265701955771059 | 0.5604144832553903 | 0.10686 | -0.06371 | -0.0349511764 |
| ***ZNF294*** | 0.843771624355496 | 0.7523678473803191 | 0.48573275021457296 | -0.01665 | 0.01150 | -0.0361461683 |

[a]The GELS, adapted from Ballhausen et al. (2023), are based on MHC ligand prediction and the prevalence of the respective HLA allele in European Caucasians, with a conservative estimate of predicted HLA-binding probability of $p_{binding}$ = 50%.
[b]OLLs were adapted from Witt et al. (2023).

*B2M, B2-microglobulin; cMS, coding microsatellite; CRC, colorectal cancer; m, m1-frameshift mutation frequency; OLL, overall ligand likelihood*
